# Cognition-Associated Protein Structural Changes in a Rat Model of Aging are Related to Reduced Refolding Capacity

**DOI:** 10.1101/2024.09.20.614172

**Authors:** Haley E. Tarbox, Audrey Branch, Stephen D. Fried

## Abstract

Cognitive decline during aging represents a major societal burden, causing both personal and economic hardship in an increasingly aging population. There are a few well-known proteins that can misfold and aggregate in an age-dependent manner, such as amyloid β and α-synuclein. However, many studies have found that the proteostasis network, which functions to keep proteins properly folded, is impaired with age, suggesting that there may be many more proteins that incur structural alterations with age. Here, we used limited-proteolysis mass spectrometry (LiP-MS), a structural proteomic method, to globally interrogate protein conformational changes in a rat model of cognitive aging. Specifically, we compared soluble hippocampal proteins from aged rats with preserved cognition to those from aged rats with impaired cognition. We identified several hundred proteins as having undergone cognition-associated structural changes (CASCs). We report that CASC proteins are substantially more likely to be nonrefoldable than non-CASC proteins, meaning they typically cannot spontaneously refold to their native conformations after being chemically denatured. The potentially cofounding variable of post-translational modifications is systematically addressed, and we find that oxidation and phosphorylation cannot significantly explain the limited proteolysis signal. These findings suggest that noncovalent, conformational alterations may be general features in cognitive decline, and more broadly, that proteins need not form amyloids for their misfolded states to be relevant to age-related deterioration in cognitive abilities.

**TEASER:** Up to 10% of rat hippocampal proteins can undergo structural changes that associate with age-related decline in spatial learning.

## INTRODUCTION

Cognitive decline in several memory domains is a common feature of aging, even in the absence of neurodegenerative disease. However, cognitive decline is not inevitable; some individuals retain cognitive abilities on par with young adults well into their later decades of life. Identifying molecular features that are retained by these cognitively resilient individuals or those that are altered in impaired subjects can provide opportunities for prevention or treatment of age-related cognitive decline. Individual variability in cognitive aging is not unique to humans. The use of animal models, such as outbred rats, has established that cognitive integrity at older ages is due to retaining functional circuit computations within an aging brain context. For example, in the well-studied medial temporal lobe (MTL) episodic memory circuit, hippocampal subfields do not exhibit overt cell loss in aged subjects with MTL-related memory impairments (*1*), and neuronal connections between hippocampal subfields are generally preserved (*2*, *3*). However, research has provided ample evidence for functional alterations in the hippocampus that are closely tied to age-related cognitive decline (*4–6*), which are linked to cellular and molecular changes (*7–9*).

In the context of a largely structurally preserved MTL, age-related changes in the proteome emerge as a potential mediator of cognitive circuit dysfunction leading to memory impairment. While protein aggregation plays a causal role in several late-life neurodegenerative diseases which impair cognition, this is largely related to the neurotoxic effects of aggregates. Amyloids possess distinctive and stable structures, which make them visible through many methods on several length scales (x-ray crystallography to histological staining). It is notable that very few proteins have been shown to misfold and/or aggregate in this dramatic way. However, proteins can misfold in an age-dependent or cognition-relevant manner but do so in a way that retains them in a soluble misfolded (*10*) and/or oligomeric (*11*, *12*) form, which would be challenging to detect with most techniques.

The proteostasis network is critical for maintaining proper protein structure *in vivo*, and comprises the processes and factors responsible for maintaining the proper balance of protein synthesis, folding, maintenance, and degradation (*13*). Loss of correctly-functioning cellular proteostasis has been shown to arise with age (*14–16*), and combination of chaperone dysfunction, turnover impairment, and PTM accumulation that occur with age creates the ingredients for proteome-wide protein structural changes which could cause changes in function. There is already evidence for specific proteins adopting soluble misfolded forms during aging. For example, altered conformations of phosphoglycerate kinase (PGK) and enolase were found in aged rat brains (*17*) and aged nematodes (*18*), respectively, among other examples (*19*). However, these studies have been limited to single proteins, and age-related protein structural changes have yet to be widely studied on a proteome scale.

Limited-proteolysis mass spectrometry (LiP-MS) is a structural proteomic technique that is able to distinguish subtle structural changes across many proteins in a complex mixture (Fig. 1B) (*20*, *21*). These structural changes are detected by comparing differences in proteolysis patterns, and can arise from conformational changes, differences in ligand binding, post-translational modification, protein-protein interactions or oligomeric state. LiP-MS has already been applied in an aging context to study protein structural changes in cerebrospinal fluid (CSF) from young and aged mice (*22*) and in young and aged yeast extracts (*23*).

**Figure 1.**
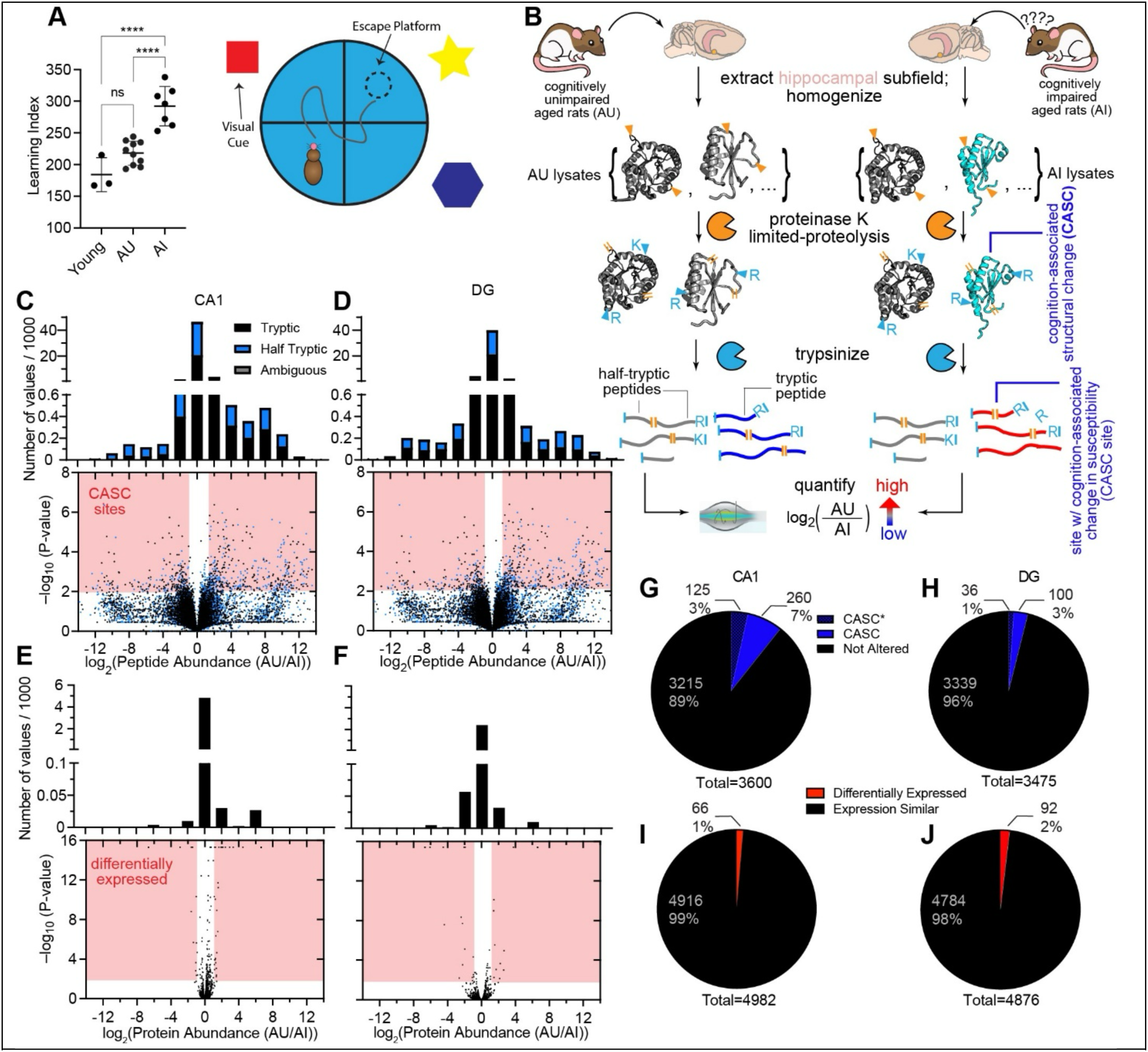
Measuring Protein Conformational Changes Associated with Age-Associated Changes in Cognition with LiP-MS. (A) The learning indexes (*31*) as assessed with the Morris water maze test (shown schematically, at right) on the twenty Long-Evans rats used in this study. Aged (24 mo) unimpaired (AU) rats have learning indexes comparable to young (∼6 mo) rats, while aged impaired (AI) rats have a significantly higher learning index than young and AU rats (****: p < 0.0001 by ANOVA with Brown-Forsythe correction and follow-up with Tukey’s multiple comparisons test). (B) Experimental scheme for limited proteolysis mass spectrometry (LiP-MS) to assess cognition-associated structural changes. Proteins in hippocampal subfields from AU and AI rats are assessed for structural differences by performing pulse proteolysis with proteinase K under native conditions followed by complete digest with trypsin under denaturing conditions. The proteolysis pattern is measured with liquid chromatography mass spectrometry (LC-MS) and label-free quantification. (C, D) Volcano plots showing changes in peptide abundance of tryptic (black) and half-tryptic (blue) peptides in the CA1 subfield (C) and dentate gyrus subfield (DG, D) between AU and AI rats. Dots that fall in the regions in red are deemed significant based on effect size (>2-fold) and p < 0.01 by t-test with Welch’s correction for unequal population variance. Histograms show number of peptides possessing abundance ratios in the various ranges. (E, F) Volcano plots showing changes in protein abundance in the CA1 subfield (E) and the DG subfield (F) between AU and AI rats. Dots that fall in the regions in red are deemed significant based on effect size (>2-fold) and p < 0.01 by t-test with Welch’s correction for unequal population variance. Histograms show number of proteins possessing abundance ratios in the various ranges. (G, H) Number of proteins with cognition-associated structural changes (CASC) in the CA1 subfield (G) and the DG subfield (H). CASC proteins have two or more peptides with significant changes. CASC* denotes proteins in which one (or more) of the significant peptides could be assigned to multiple isoforms. (I, J) Number of proteins with different measured abundance in AU vs AI subjects in the CA1 subfield (I) or DG subfield (J).

In this study, we utilize LiP-MS to interrogate the structural neuroproteome of the hippocampus in an aging rat model. In this model, genetically variable, outbred aged rats are categorized as being <u>a</u>ged and cognitively <u>u</u>nimpaired (AU) or <u>a</u>ged and cognitively impaired (AI) based on performance in the Morris water maze (MWM) test (Fig. 1A) (*24*). AI rats exhibit subfield-specific alterations in medial temporal lobe (MTL) physiology and function, which mirror those observed in aged humans with memory impairments and have successfully guided human clinical studies (*25*, *26*). AU rats execute MTL-dependent tasks on par with young animals, while exhibiting age-adaptive molecular (*7*, *27*), cellular (*28*, *29*), and circuit mechanisms (*2*, *4*, *30*). Thus, cognitively characterized aged Long Evans rats can serve as a powerful model system for assessing how individual differences in protein structures may contribute to or prevent age-related cognitive decline. We reasoned the subfields of the hippocampus in this model could be particularly relevant contexts for assessing changes in protein folding, given the extensive characterization of this circuit in both young and aged subjects. Moreover, we focus on a comparison between two aged populations in this study to isolate molecular factors associated with a cognitive phenotype, in contrast to comparing young and old populations which could convolve other factors.

In this study, we assess protein structural changes in the rat hippocampus, covering 6276 distinct proteins, and we report evidence that many (466) proteins can possess soluble structural changes that different between cognitive phenotypes. We identified proteins with cognition-associated structural changes (CASCs) by comparing the proteolysis patterns between these cohorts. A number of these proteins have been previously implicated in memory, aging, and synaptic health, whereas many others represent potentially novel targets for further study or therapeutics. Perhaps most strikingly, we discover that CASC proteins are generally nonrefoldable, that is, they often are unable to spontaneously return to their native conformations following global denaturation. Nonrefoldability explains more of the cognition-associated changes than oxidation or phosphorylation levels, suggesting that noncovalent, conformational alterations may be a general feature underlying cognitive decline. Because these structural changes are observed in a background wherein all animals are of the same age, this study reports directly on proteins that may be relevant to cognition. More broadly, our findings suggests that many proteins may misfold in more subtle ways, and unlike amyloids, remain soluble, yet still be relevant to deterioration in cognitive abilities.

## RESULTS

### Identification of Cognition-Associated Structural Changes Proteome-Wide in the Rat Hippocampus

Long-Evans rats were aged to 24 months in a colony with a controlled diet and environment. Young animals in this population (6 mo) exhibit proficient spatial memory as evidenced by their performance on the Morris water maze test, quantified by the learning index scaled described previously (*31*). Aged rats, on the other hand, exhibit a wider spectrum in their capacity to perform these tasks effectively (Fig. 1A) with some individuals exhibiting impaired spatial learning abilities relative to young subjects. These aged-impaired (AI) rats have been shown to have MTL circuit dysfunction including subfield-specific alterations in synaptic density (*2*), disruption of excitatory/inhibitory (E/I) balance (*4*, *25*), and baseline and learning-related transcription (*7*). Our study included 7 AI rats, 10 AU, and 3 young rats. Approximately 2 weeks after MWM cognitive assessment, hippocampal tissue was extracted and dissected into DG, CA3, and CA1 subfields. We focused on the hippocampus because it is closely associated with spatial learning and memory in rats (*32*, *33*).

The subfields were Dounce homogenized into a native buffer without detergents, cellular debris and membranes were depleted by centrifugation, and the soluble fractions were subjected to limited proteolysis (LiP) with proteinase K (PK, 1:100 w/w) for 1 min (Fig. 1B). This treatment enables cleavage to occur only at solvent-accessible locations within proteins and thereby encodes structural information about the conformational ensemble of each protein into cut-sites. To determine the locations of these cut-sites, the samples are trypsinized, and the resulting peptides identified through liquid chromatography tandem mass spectrometry (LC-MS/MS). In particular, sequenced peptides for which one of the termini can be assigned to a PK cleavage (orange lines in Fig. 1B, called half-tryptic peptides) because it does not follow trypsin’s cleavage rules specify a site of solvent accessibility in the parental protein structure. Changes in abundance of the resulting half-tryptic (and tryptic) peptides can be interpreted in terms of changes in proteolytic susceptibility at the corresponding sites. Here, we assess structural changes in hippocampal proteins between young (Y), AU, and AI cohorts by performing label free quantification (LFQ) on the 20 (or 19) samples, and studies of this kind were performed on each of these three hippocampal subfields separately. In general, slightly fewer peptides/proteins are identified in LiP runs (Fig. S1A-F), owing to the expanded search space and the treatment degrades some protein regions, but overall similar numbers of peptides/proteins were identified across runs, subfields, and cohorts. We also performed a parallel “trypsin-only” set of samples in which the limited proteolysis step was withheld; the resulting quantifications provide a means to assess protein abundance differences between AU and AI rats and to normalize the peptide abundance ratios from the LiP samples with overall protein-level abundance ratios. Importantly, a relatively small fraction of the identified peptides in the trypsin-only control samples were of the half-tryptic variety (10.1 ± 1.1 %), in contrast to that of LiP samples (49.3 ± 6.2 %; Fig. S1G-I). This suggests that most of the half-tryptic peptides we sequenced correspond specifically to cleavage from structural probing by PK (and not by sample degradation or endogenous protein decay) and was possible due to an amended LiP-MS protocol in which extracts were treated with a specific cocktail of protease inhibitors to mitigate degradation in the lysate without interfering with PK (see SI *Methods*).

In Figure 1C-D, we show normalized peptide-level volcano plots summarizing the results from these experiments (see SI *Methods*) in the CA1 and DG subfields; on these diagrams, each point represents a confidently identified and quantified peptide. We quantified 54,614 and 49,058 peptides respectively, of which 1,578 (2.9%) and 863 (1.8%) were above our thresholds (2-fold effect size, P-values < 0.01 by two-tailed t-test with Welch’s correction) to be considered significantly enriched in either AU or AI hippocampi. Next, we assigned these peptides to their parental proteins, and if we detected two or more uniquely-mappable peptides with significantly altered proteolytic susceptibility, the protein is labeled a CASC protein (cognition-associated structural change). By this metric, 260 proteins are CASC in CA1, and 100 are CASC in DG (Fig. 1G-H), representing 7% and 3% of the total proteins assessed. We moreover verified that LiP-MS quantifications are highly reproducible on hippocampal extracts in a separate study (Fig. S2) in which we compared difference in measured peptide levels across different injections (measuring instrumental variability), different LiP sample preparations (measuring variability from the LiP protocol), and different subjects (measuring biological variability).

Interestingly, we detected markedly fewer differences in the CA3 subfield between the AU and AI cohorts (Fig. S1J-M). While one explanation of these results could be that the CA3 preparations had higher within-sample variation, we found instead that the within-sample variation for CA3 is comparable to the variation observed for the CA1 preparations (Fig. S3D-F, J-L). This suggests that the relative lack of CASC proteins in CA3 is unlikely to be a result of technical or LiP-related reproducibility issues. Additionally, these observations are qualitatively consistent with prior work suggesting the CA3 subfield of the hippocampus may possess some relative protection to stresses related to aging, such as oxidative stress, compared to the CA1 subfield (*34*, *35*).

It is notable to point out that at the protein-level, there are comparatively fewer differences between the AU and AI cohorts (Fig. 1E-F). In CA1, 66 proteins are differently expressed, and in DG, 92 proteins are differentially expressed (Fig. 1I-J). Hence, using structural proteomic methods can report differences on more proteins than a “standard” omics approach, which only reports on protein abundances, when comparing two wild-type populations which vary in their cognitive phenotype.

### Proteins with Cognition-Associated Structural Changes Are Less Capable of Spontaneously Refolding Following Denaturation *in vitro*

Previous work of ours (*36*, *37*), has applied LiP-MS to address what might outwardly appear to be an unrelated question: Which proteins are incapable of spontaneously refolding to their native structures following complete denaturation? In these experiments, an extract is subjected to a global unfolding-refolding cycle (Fig. 2A) in which proteins are first incubated overnight in 6 M guanidinium chloride (GdmCl) and 10 mM dithiothreitol (DTT), and then diluted 100-fold to initiate refolding. Following several refolding times, the proteome-wide refolding reactions are probed by LiP and the resulting proteolysis pattern compared to that of native (unperturbed) samples that were not subjected to an unfolding-refolding cycle. We consider these experiments as a probe of a protein’s spontaneous (unassisted) refolding capacity because the chaperone and ATP concentrations in the diluted lysates are too low to play a significant role unless they are supplemented to cellular-like levels (*38*). Our previous work demonstrated that after 5 min (a time that allows many proteins sufficient time to refold but prior to significant degradation), 60% of *E. coli* proteins can refold (*36*), and 76% of yeast proteins can refold (*37*). We performed similar global refolding assays on hippocampal extracts from young rats and found an overall refolding rate after 5 min of refolding (74%) more similar to yeast (Fig. 2B), hence we continued to use this timepoint as the reference condition (Fig. S4C). As previously, to be labeled nonrefolding, a protein needed to possess at least two sites with altered susceptibility (2-fold effect size, P-values < 0.01 by two-tailed t-test with Welch’s correction) in the refolded form. At dilute concentrations of 0.1 mg/ml, global refolding reactions on the hippocampal extract could proceed without any detectable level of precipitation (Fig. S4A), and the refolded proteins were, on the whole, more susceptible to limited proteolysis (Fig. S4B), suggesting the formation of soluble misfolded conformations. The abundance of LiP peptides in refolding reactions from biological replicates were reproducible (Fig. S3A-C), and the 5 min refolding reactions possess a median coefficient of variation of 14% (Fig. S3B), and other quality controls were acceptable, such as consistent peptide/protein IDs across replicates, low incidence (13.4 ± 0.4%) of half-tryptic peptides in trypsin-only controls and an expected frequency (51.7 ± 4.2%) of half-tryptic peptides in LiP samples (Fig. S4D-F). It is notable that the variation across refolding reactions is much lower than the biological variation we encountered in the aging study (range of medians from refolding study: 0.13-0.17; range of medians from aging study: 0.26-0.67, Fig. S3A-L).

**Figure 2.**
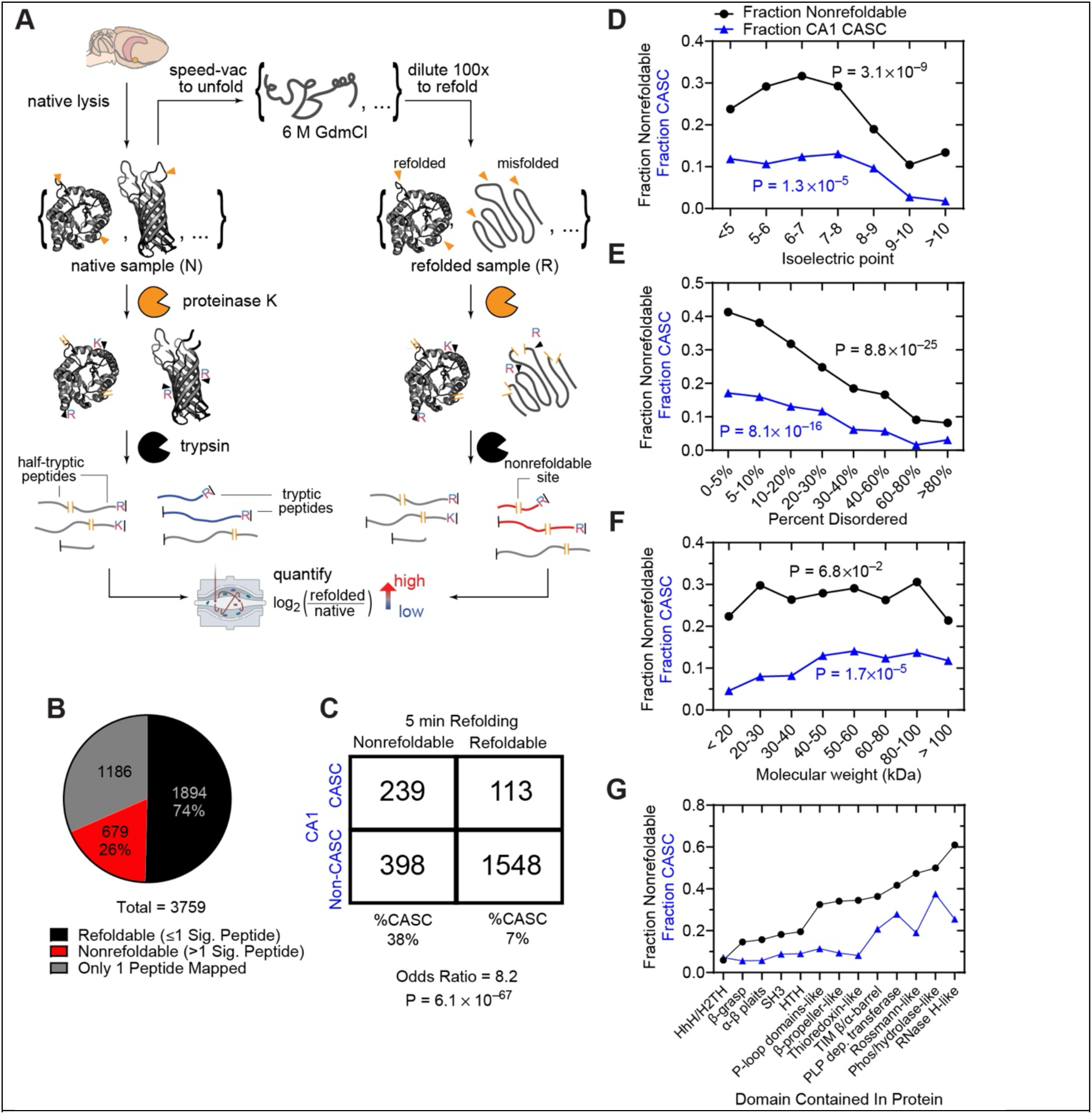
Similarities Between Nonrefoldable Proteins and Proteins with Cognition-Associated Structural Changes. (A) Experimental scheme for limited proteolysis mass spectrometry (LiP-MS) to assess refoldability. Proteins in the young rat hippocampus are unfolded by incubation in 6 M guanidinium chloride (GdmCl) and are refolded by 100-fold dilution into a native buffer. The structures of the native proteins and their refolded forms are compared by performing pulse proteolysis with proteinase K followed by complete digest with trypsin, and the proteolysis profiles are measured. (B) Number of refoldable (black) and nonrefoldable (red) proteins in the hippocampus. Nonrefoldable proteins have two or more peptides with significant changes following refolding. Proteins with only 1 peptide mapped are discounted (gray). (C) Contingency table comparing refoldability status in the global refolding studies and CASC status in the cognition studies. Refolding after 5 min and CASC status in the CA1 were used as the reference conditions, respectively. Shown below the table are the marginal frequencies of CASC status among (non)refoldable proteins, the odds ratio, and the p-value against the null hypothesis of independence according to Fisher’s exact test. (D-G) Plots provide the fraction of proteins within a given category that are nonrefoldable after being allowed 5 min to refold from denaturant (black) or that are CASC in hippocampal CA1 (blue) as a function of: (D) protein isoelectric point (pI), (E) percent disorder according to Metapredict (*39*), and (F) molecular weight. Panel G assesses proteins based on whether they contain a domain of a given topology (based on ECOD (*40*)) though these categories are not mutually exclusive since some proteins contain multiple domains. P-values (according to chi-square test) against the null hypothesis that nonrefoldability/CASC status is independent of the categorical variable in question are provided.

To compare the outcomes of these two experiments, we considered each protein whose refoldability and CASC status (in CA1) could be simultaneously assessed. The results in the contingency table (Fig. 2C) show that these statuses are highly associated (P-value < 10^-66^ by Fisher’s exact test). For instance, refoldable proteins have only a 7% likelihood of being CASC, but nonrefoldable proteins have a 38% (5.4-fold greater) likelihood to be CASC.

Some structural parameters, such as isoelectric point and disorder content, display similar trends for CASC and nonrefoldable proteins. For example, proteins with an isoelectric point between 6 and 8 are most likely to be CASC or nonrefoldable, while proteins with an isoelectric point above 9 are among the least likely (Fig. 2D). Similarly, as the protein’s disorder content (as calculated by Metapredict (*39*)) increases, the likelihood of a protein being classified as CASC or nonrefoldable generally decreases (Fig. 2E).

One limitation of our metric to ascribe nonrefoldable or CASC status (2 or more sites on a protein with altered susceptibility) is that proteins with more peptides mapped to them are more likely to have altered peptides. One way this bias can be evaluated is by assessing all the peptide sites without first assigning them to proteins (*36*). For instance, we find that sites from proteins whose isoelectric point are between 6 and 8 have the highest propensity to be structurally altered following unfolding/refolding, implying that the high incidence of nonrefoldability amongst this class of proteins is not due to coverage bias, as this trend recapitulates what was seen on the protein level (Fig. S5A). Similarly, we find that peptides from proteins with isoelectric points between 6 and 8 also have the highest propensity to be CASC sites (altered susceptibility between AU and AI) (Fig. S5B). Similarly, the frequency of altered peptides decreases with increasing disorder content on both the protein and peptide level for nonrefoldability and CASC status (Fig. S5C-D).

Analogously to what we previously observed in yeast (*37*), molecular weight is not a major determinant of refoldability (Figs. 2F, S5E). While it may appear that molecular weight does slightly positively correlate with CASC (Fig. 2F), this effect is not robust as it is not recapitulated at the peptide level (Fig. S5F).

One of the more striking ways that nonrefoldability and CASC statuses cohere with one another is by noting which types of domain topologies are contained in these proteins. Amongst proteins we observe in the rat hippocampus, all-α and all-β domains (such as helix-turn-helix (HTH), helix-hairpin-helix (HhH), SH3 folds, β-grasp (the ubiquitin-like domain)) typically refold efficiently, but larger α/β domains – particularly those involved in metabolic processes, such as TIM barrels, and PLP-dependent transferases – have low refolding capacity (Fig. 2G). This mimics what we observed in a recent study investigating refoldability in the yeast proteome (*37*). Moreover, the same types of domains that exhibit higher nonrefoldability also tended to have high propensity to become structurally perturbed in a cognition-dependent manner (Fig. 2G, compare blue and black traces). For example, over 40% of proteins that contain a phosphorylase/hydrolase-like domain, or a pyridoxal phosphate (PLP) dependent transferase domain, are nonrefoldable. Over 27% and 37%, respectively, of proteins containing these same domains are CASC proteins. On the other hand, proteins containing domains such as alpha-beta plaits or SH3 are overwhelmingly refoldable (<20% nonrefoldable) and are also unlikely to be CASC proteins (<10%). This trend appears robust based on similar nonrefolding/CASC propensities being observed at both the protein and peptide levels (Fig. S5G-H).

### Critical Proteins Emerge as CASC Proteins

While looking at bulk trends of the characteristics of proteins is instructive in getting a sense of what types of proteins are likely to possess structural differences between the AU and AI populations, we investigated several proteins in greater detail. For example, DDX39B (Uniprot accession Q63413) is a DEAD-box family RNA helicase that also has RNA-binding activity and acts as a splice factor (*41*). Another DEAD-box RNA helicase involved in splicing activity, DDX5, has recently been shown in killifish to form aggregates with age (*42*). We modeled sites of nonrefoldability (shown in black), CASC (shown in light blue), or sites that appeared as significant in both experiments (dark blue) onto DDX39B’s AlphaFold2 structure(*43*) (Fig. 3B). For half-tryptic peptides, the residue associated with the PK cut is shown, whereas for tryptic peptides, the middle residue for the sequenced tryptic fragment is shown. Two sites are identical between the nonrefoldability and CASC data sets, and the remaining CASC site is near these two sites. The N-terminal portion of the 2^nd^ P-loop domain in the protein can become structurally altered both during natural processes in the hippocampus as well as during a misfolding pathway that occurs after dilution from denaturant *in vitro*.

**Figure 3.**
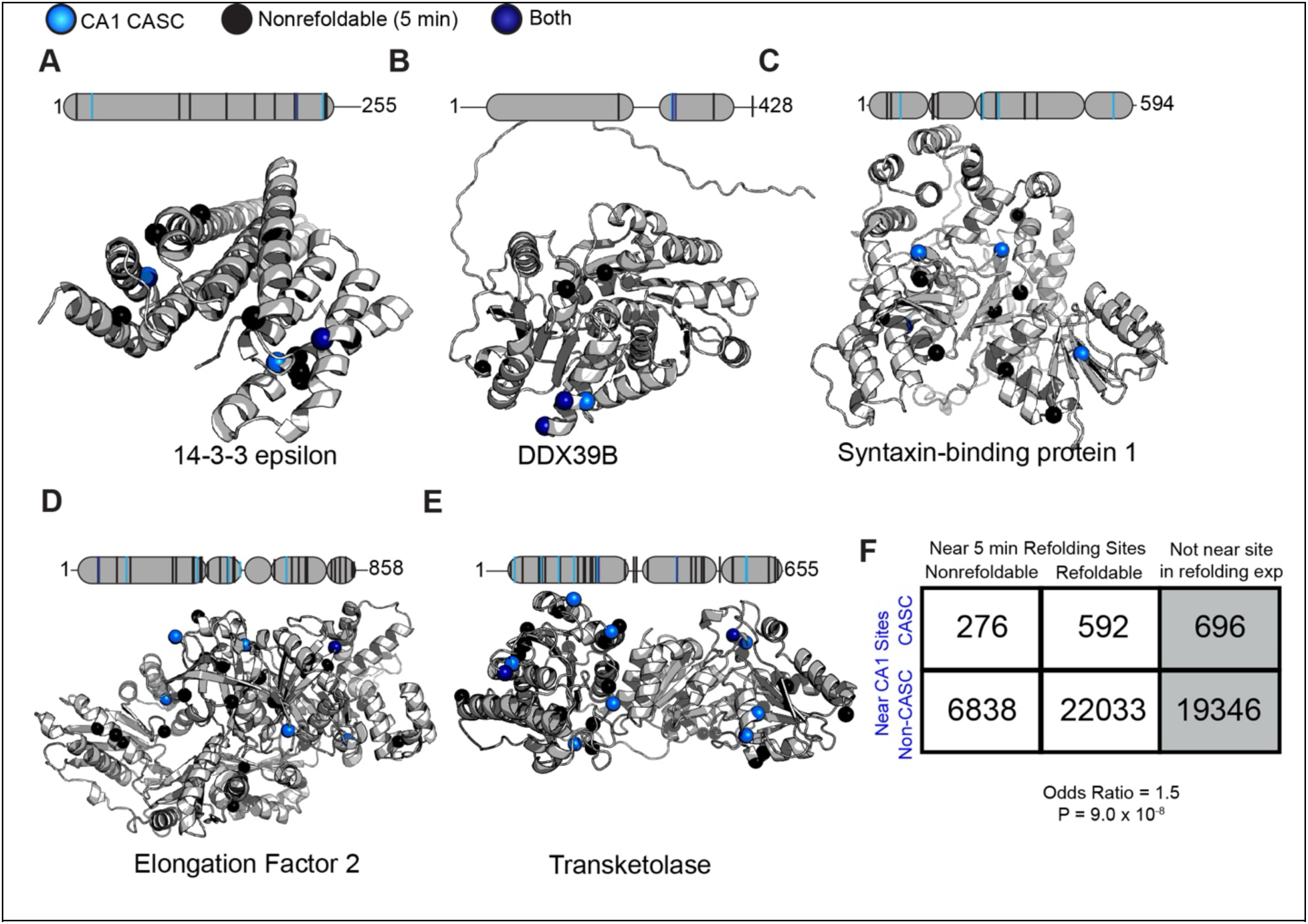
Colocalization Between Nonrefoldable Sites and Sites with Cognition-Associated Structural Change. (A-E) AlphaFold2 structural models displayed for (A) 14-3-3 protein epsilon (P62260), (B) DDX39B (Q63413), (C) Syntaxin-binding protein 1 (P61765), (D) eukaryotic elongation factor 2 (P05197), and (E) Transketolase 1 (P50137) showing the locations of residues that are structurally altered (based on changes in proteolytic susceptibility) after 5 min of refolding from denaturant in global refolding assays (black spheres) or structurally altered in CA1 between AU and AI rats (light blue spheres). Residues that are structurally altered in both experiments are shown as dark blue spheres. Sequence diagrams (above each structure) show the positions of these residues with respect to the domain boundaries. Indicated positions represent cut-sites for half-tryptic peptides and midpoint residue for tryptic peptides. (F) Contingency table showing the number of sites that are (non)refoldable in global refolding assays and/or structurally altered in a cognition-associated manner. To be included, a site identified in the cognition experiment had to be within five residues on either side of a site identified in the refolding experiment. Gray represents sites excluded from the analysis because they were not identified in the refolding experiment. P = 9.0 × 10^-8^ by Fisher’s exact test.

We modeled several other proteins, including 14-3-3 protein epsilon (P62260), syntaxin-binding protein 1 (P61765), elongation factor 2 (P05197), and transketolase (P50137), all using their AlphaFold2-predicted structures for consistency (Fig. 3A,C-E). These five proteins identified by our study, all of which are CASC in both CA1 and DG, already have some ties to aging and cognitive decline in many cases. For example, 14-3-3 proteins are involved in regulating levels of hippocampal postsynaptic NMDA receptors (*44*), which are important for synaptic plasticity. Additionally, lower NMDA receptor activity has been suggested to lead to neuronal hyperactivation (*45*), which is a phenotype seen in AI rats (*46*). 14-3-3 epsilon specifically has been implicated in neurogenesis in DG and identified as a potential protein of interest for cognitive decline in aging (*47*). 14-3-3 proteins in general have also been identified as having potential molecular chaperone-like activities (*48*). Syntaxin-binding protein 1 is an essential protein involved in synaptic vesicle docking and neurotransmission (*49*), and it has been investigated previously for potential to aggregate in brain tissue (*50*). Elongation factor 2 is essential for protein synthesis, and lower activity has been demonstrated with age (*51*), suggesting a possible age-related structural change. A decrease in transketolase (a key enzyme in the pentose phosphate pathway) activity in human blood has been observed in an age-dependent manner (*52*).

These models also demonstrate that nonrefoldable sites and CASC sites are sometimes colocalized, in particular for DDX39B, 14-3-3 epsilon, and transketolase (Figs. 3A-B, E). On a more global scale, we performed a proximity analysis to determine if CASC sites (from the aging/cognition study) and nonrefoldable sites (from the refolding study) were near one another (within 5 residues on either side). CASC sites indeed tend to be found near nonrefoldable sites (31.8%), which occurs at a somewhat higher frequency than non-CASC sites being found near nonrefoldable sites (23.7%; Fig. 3F).

Although the proximity effect is statistically significant (P-value < 10^-7^ by Fisher’s exact test), we note that the effect-size for this association (odds ratio = 1.5) is less pronounced than what we previously noted when considering entire proteins instead of peptides (Fig. 2C). The interpretation of these findings is that while nonrefoldable proteins have a greater susceptibility to become structurally altered in cognitive impairment, the precise sites within proteins that misfold in protein refolding experiments do not always closely map onto the sites associated with cognitive impairment. For instance, in syntaxin-binding protein 1 and elongation factor 2, the sites altered during refolding and cognitive decline are more diffuse and are not as close to one another.

Since structure is often related to function, we wanted to investigate if our CASC proteins demonstrated a difference in activity between AU and AI hippocampi. We chose two CASC proteins, phosphoglycerate kinase (PGK) and transketolase. PGK is a glycolytic enzyme, and transketolase is involved in the pentose phosphate pathway

(*53*). We performed activity assays (phosphorylation of 3-phosphoglycerate for PGK and two-carbon transfer from ribose-5-phosphate to xylulose-5-phosphate, see SI *Methods*) with three AU and three AI hippocampal lysates and observe Michaelis-Menten kinetics for both enzymes (Fig. S6A-B); however, we did not observe significant differences in the Michaelis constant (K_m_) or in the maximum velocity (V_max_) for either PGK or transketolase (Fig. S6C-F). Western blots showed there was no significant differences in expression level for PGK and transketolase between AU and AI hippocampi (Fig. S6G-J). While we did not see any activity differences between these two proteins, these are two representative examples out of several hundred CASC proteins, wherein a closer coupling between structural changes and changes in activity might be observed in future studies.

### Post-Translational Modifications Do Not Explain Cognition-Associated Structural Changes

Post-translational modifications are commonly associated with aged proteins (*54*) and can affect also structure and function (*55*). Moreover, PTMs could potentially be confounding factors in our limited-proteolysis experiment (Fig. 4A). First, the presence of a PTM could theoretically change the susceptibility of PK at a site near the modification (Fig. 4A, case *iii*), which could change the levels of half-tryptic peptide formed, resulting in a site being categorized as a CASC site, thereby acting as an indirect reporter of the change in PTM state between AU and AI subjects.

**Figure 4.**
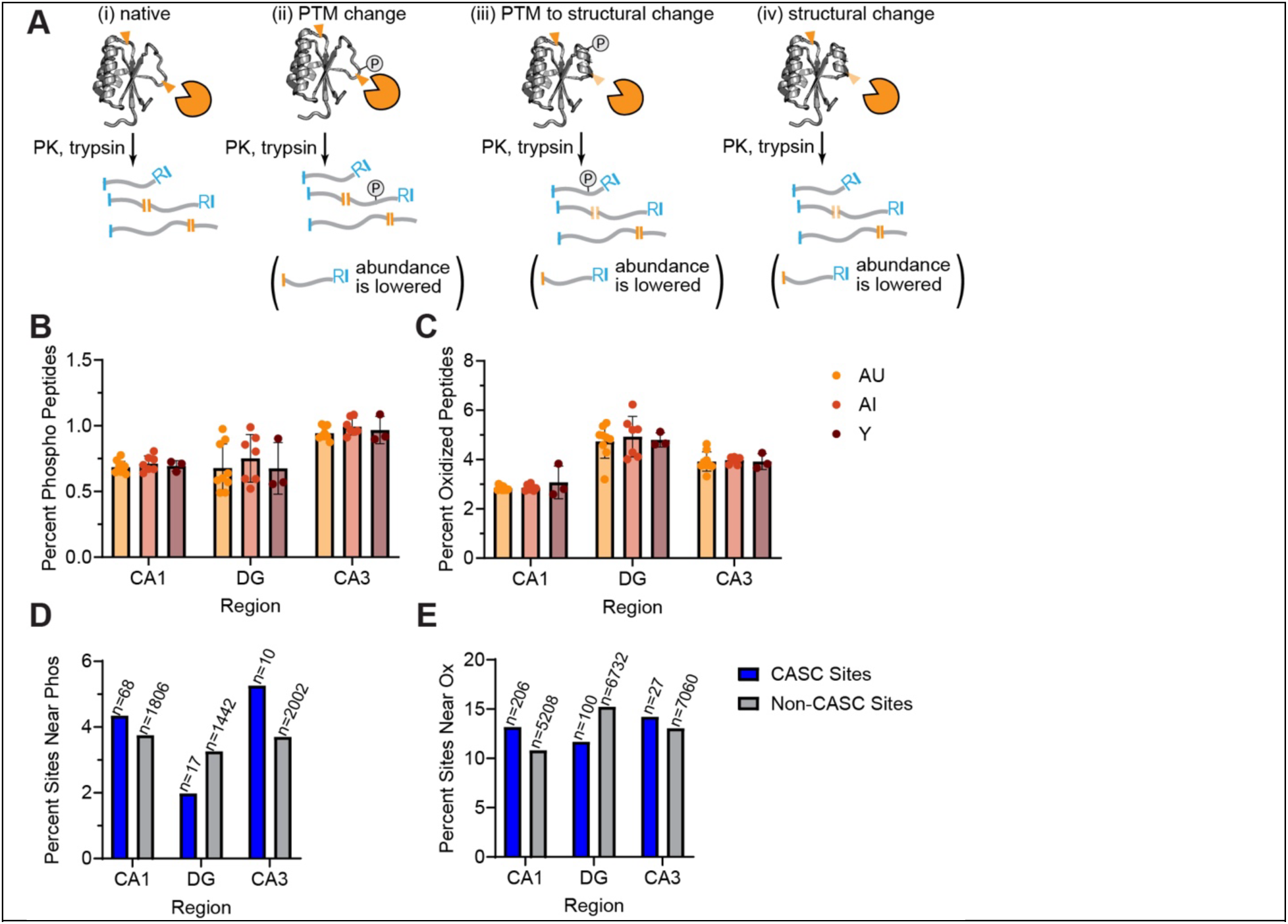
Post-translational Modifications Do Not Explain the Preponderance of the Cognition-based Changes in Proteolysis. (A) Scheme showing three scenarios (labeled cases ii, iii, and iv) for how the abundance of a half-tryptic peptide might be changed during limited proteolysis relative to a reference (case i). Case ii indicates a PTM directly changing a peptide abundance, since the mass of the peptide has changed with the modification; case iii indicates a PTM indirectly changing peptide abundance via an associated structural change; case iv indicates a change in proteolytic susceptibility only due to a structural (non-covalent) change. (B, C) Bar charts showing the percent of total identified peptides (filtered to an FDR of 1%) that contain a phosphorylation (at Ser, Thr, or Tyr; panel B) or oxidation (at Met, Phe, Tyr, Trp; panel C) from trypsin-digested peptides from hippocampal extracts associated with three subfields and three rat cohorts (young (Y), aged-impaired (AI), aged-unimpaired (AU)). Dots represent individual biological replicates, bar heights represent averages, error bars represent std. dev. (D, E) Bar charts showing the percent of peptides with significant cognition-associated changes (blue) and with no cognition-associated changes (gray) that are proximal to any detected phosphorylation site (panel D) or oxidation site (panel E). Proximity is within 15 residues on either side of the cut-sites for half-tryptic peptides, and within 15 residues on either side of the midpoint residue for tryptic peptides. Numbers above bars represent the total number of CASC and non-CASC sites within the given hippocampal region proximal to the PTM.

Additionally, the mass and ionizability of a peptide containing a PTM compared to a non-modified peptide are different; hence modification will decrease the abundance of a non-modified peptide, but it would be incorrect to ascribe such a peptide abundance difference to a structural change when it is simply due to the existence of the PTM (Fig. 4A, case *ii*). We performed several analyses to test whether these effects could explain the cognition-associated changes in proteolytic susceptibility at hundreds of sites we identified. First, we re-searched all of the non-LiP raw data for phosphorylation modifications (+79.966 Da on Ser, Thr, and Tyr) and for oxidation modifications (+15.995 Da on Met, Phe, Tyr, and Trp). We could not find a meaningful difference in the frequency of peptide phosphorylated between AU, AI and young rats within any region (CA1, DG, nor CA3) (Fig. 4B). We also could not see a difference in the frequency of peptides with any oxidation modifications (Fig. 4C). Therefore, varying amounts of modifications between the groups (i.e., AU and AI) do not confound our LiP experiment, at least at the overall statistical level.

Next, we asked if these modifications tended to be localized near PK cut-sites, where they could potentially cause differences in PK susceptibility, which would be detected as a “significant” peptide by LiP-MS. Specifically, we looked at PK cut-sites (or centers of full-tryptic peptides) on all CASC sites (and all non-CASC sites), and asked if there were any detected phosphorylation or oxidation modifications located to within 15 amino acids of the site. In general, we find that CASC sites are not close to PTMs, with 2-5% of CASC sites being close to detected phosphorylations and 10-15% of CASC sites being close to detected oxidations (Fig. 4D-E, blue bars). On the other hand, we do find a slight enrichment for CASC sites to be close to PTMs compared to non-CASC sites in the CA1 and CA3 subfields of the hippocampus (Fig. 4D-E, compare blue to gray bars), though there is not a significant association between these categories according to Fisher’s exact test for the CA1 and CA3 subfields for the phosphorylation analysis, nor for the CA3 subfield in the oxidation analysis. In DG, the effect is reversed (CASC sites are less likely to be next to modifications than non-CASC sites). Hence, these findings are consistent with the notion that the majority of the differences in PK susceptibility that we encounter in this study are explained by differences in protein structure than they are by differences in PTM status (case *iv* in Fig. 4A). Moreover, these data show that PTMs do not overwhelmingly explain our CASC site identifications, as PTMs occur at similar frequencies both near CASC and non-CASC sites, and there is no major difference in PTM frequencies between AU, AI, and young within each hippocampal subfield.

### Gene Ontology Analysis

Using ShinyGO (*56*), we investigated gene ontology (GO) terms enriched in the set of CASC proteins, focusing on the CA1 region which provided the most examples (Fig. 1G-H). GO Biological Process analysis revealed that many metabolic terms are enriched in CASC proteins, along with several terms associated with the synapse, including “modulation of chemical synaptic transmission” and “regulation of trans-synaptic signaling” (Fig. 5A). Synapses are often affected by aging (*57*, *58*) and are critical for memory formation and long-term potentiation (*59*). For instance, calcium/calmodulin-dependent protein kinase II (CaMKII), has been assigned a role in memory formation in the hippocampus by facilitating long-term potentiation (*60–62*), and it is identified as a CASC protein in this study. Specifically, its alpha isoform localizes to the synapse (*60*), and 50% (5 out of 10) of the CASC sites we found in CaMKII are specific to the alpha isoform, with the other 5 CASC sites non-isoform-specific.

**Figure 5.**
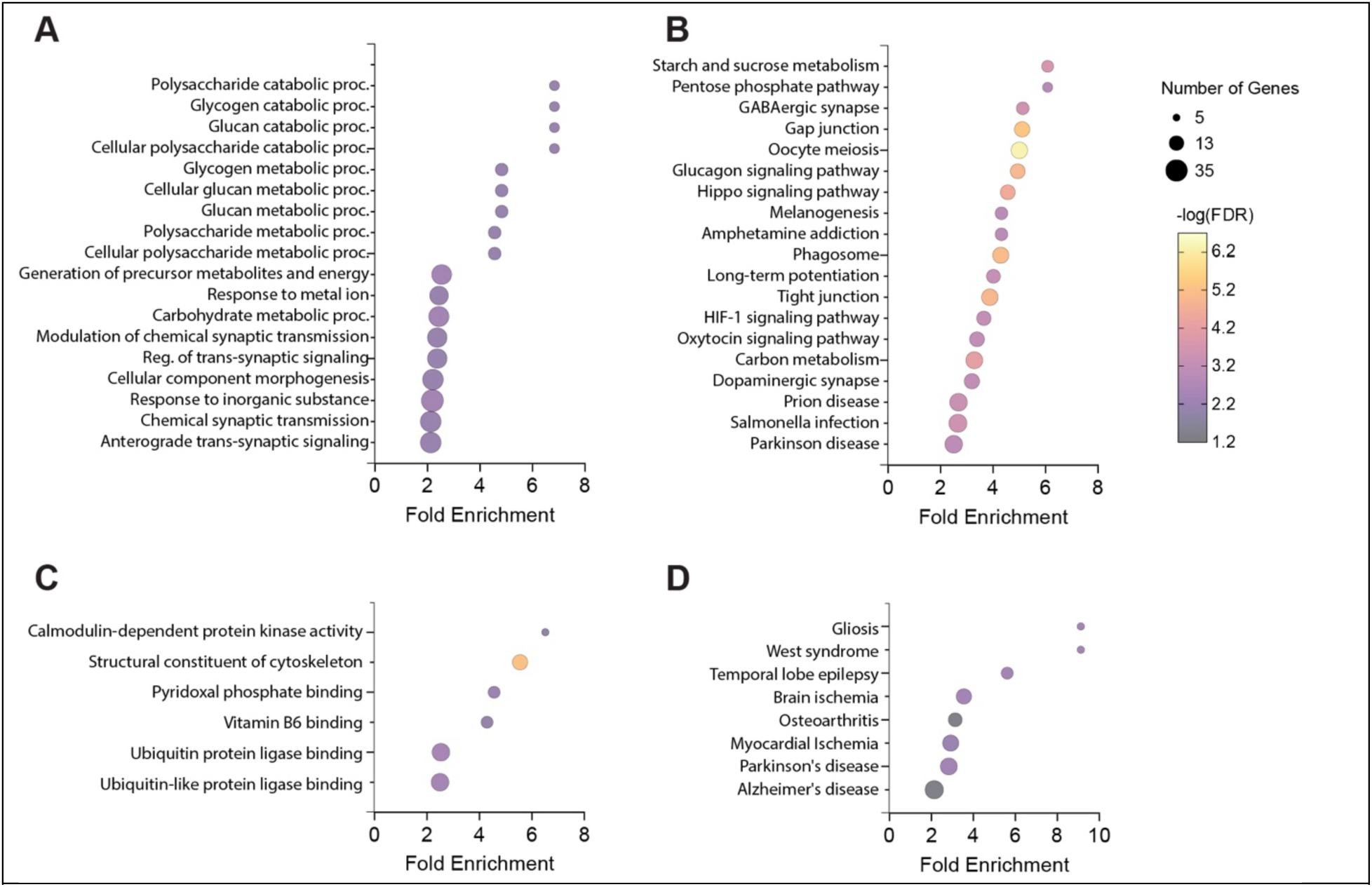
Gene Ontology Analysis of Cognition-Associated Structurally Changed Proteins. Analysis is based on the 385 CASC proteins from the CA1 subregion of the rat hippocampus, with a reference set composed of all 3600 proteins quantified with 2 or more peptides across the 17 biological replicate limited-proteolysis AU and AI CA1 samples using ShinyGO (*56*). (A) Using the biological process GO terms. (B) Using the Kyoto Encyclopedia of Genes and Genomes (KEGG) database. (C) Using the molecular function GO terms. (D) Using the genes associated with disease pathways as defined by the Rat Genome Database (RGD).

Gephyrin plays a critical scaffolding role at the synapse of inhibitory neurons (*63*), and also is a CASC protein. Some work has suggested that neuronal hyperactivity may be detrimental to the aging brain, as neuronal inhibition in *C. elegans* was shown to increase lifespan (*64*), and cognitively impaired rats have hippocampal overactivity compared to age-matched unimpaired rats (*46*). Glutamate decarboxylase 1 and 2 (Gad1 and Gad2) are also CASC proteins, which are important for the synthesis of GABA, an inhibitory neurotransmitter (*65*). Structural and functional characterization of these proteins at the synapse may help elucidate the molecular basis for neuronal overactivity and its linkage to cognitive decline.

Using the Kyoto Encyclopedia of Genes and Genomes (KEGG) database (*66*), (Fig. 5B), we see more synapse-related terms including “GABAergic synapse,” “Long term potentiation” and “dopaminergic synapse,” and many key cell-signaling pathways. A molecular function GO analysis (Fig. 5C) identifies an enrichment for ubiquitin protein ligase binding (central to the ubiquitin-proteasome system that forms a backbone of the proteostasis network), reemphasizes the high enrichment of CamKII and proteins functionally similar to it, and also calls attention to pyridoxal phosphate (PLP) binding proteins. Many PLP binding proteins utilize a particular fold (*67*, *68*) that is quite topologically complex (a 3-layer α/β/α sandwich) that we had previously observed exhibits poor refoldability in *E. coli* (*36*) and in rats (Fig. 2G). Finally, we searched for whether CASC proteins were enriched in disease-related genes from the Rat Genome Database (RGD) and found that CASC proteins are indeed enriched primarily in diseases associated with the brain (Fig. 5D).

Gene ontology analyses for the nonrefoldable proteins discovered in the refoldability study were conducted as well. Biological Process GO showed enrichment of nonrefoldable proteins in several metabolic pathways (Fig. S7A), but lacked synapse-specific terms as seen in the CASC GO analysis. KEGG analyses show nonrefoldable proteins are over-enriched in several signaling pathways (Fig. S7B). Molecular Function analyses show that ATP-dependent processes are enriched in nonrefoldable proteins (Fig. S7C) and cellular component analysis identifies a few complexes, including the proteasome, to be enriched in the nonrefoldable set (Fig. S7D). As nonrefoldability reports on protein structure and not a protein’s function in the cell, we were not surprised that it possessed fewer linkages to various GO terms.

Finally, we performed GO analyses on the non-LiP dataset assessing protein abundance differences between aged-impaired and aged-unimpaired rats (cf. Fig. 1I). We did not see any classifications significantly enriched among the 66 proteins using either the databases defined by biological processes, molecular functions, KEGG, cellular component, or RGD disease pathways. This negative result suggests that protein abundance differences in the hippocampus could not explain differences in cognitive status across this rat population, at least with the statistical power of the present study.

## DISCUSSION

Here we have employed limited-proteolysis mass spectrometry (LiP-MS) to identify proteins that vary in structure in the hippocampus of aged rats with or without cognitive impairment, which we have defined as cognition-associated structural change (CASC) proteins. CASC proteins were identified in all hippocampal subfields but were most numerous (260) in the CA1 region. Research in aging, dementia, and neurodegenerative disease has long made a connection between these diseases processes and protein misfolding; however, emphasis has historically been paid to proteins which form amyloids or other insoluble aggregates. We have focused on the soluble fraction of the hippocampal proteome, and employed a methodology that can sensitively detect subtle changes in protein structure. The results enable us to conclude that protein misfolding is perhaps a more pervasive feature in cognitive decline than previously appreciated, and that many of these misfolded forms persist as soluble species. This finding suggests that there may be new avenues for potential therapeutic targets and diagnostic biomarkers for cognitive decline than the small subset of amyloid-forming proteins frequently studied. Of course, such interventions would need to be conformation-specific, creating new opportunities and challenges.

Strictly speaking, the changes in proteolytic susceptibility that we detect (Fig. 1) could be explained by a wide range of factors, including changes in interaction partners, changes in oligomeric stage, and changes in post-translational modifications (PTMs). One of the limitations of our study is that we cannot directly identify the exact physical or functional nature of structural changes that occur in CASCs. As proteins age, they can accumulate PTMs, including enzymatic PTMs such as phosphorylation and non-enzymatic PTMs such as oxidation (*54*). These PTMs can affect the folding and the function the proteins they modify (*55*), and can also cause conformational changes, even at regions of the protein far from the PTM site itself (*69*). Our results suggest that most of the changes in proteolytic susceptibility observed in our assay can be ascribed to changes in conformation or misfolding. We were able to rule out the possibility that PTMs explain the majority of the signal because they can also be localized by mass spectrometry, and we could not find evidence for strong co-localization to CASC sites. While it is likely that some of the CASCs we identify report on changes in interaction partners or oligomeric state, the exceedingly strong overlap between CASC status and nonrefoldability (Fig. 2C) provides indirect evidence that these features arise from a common underlying etiology in a protein being complicated to fold or prone to misfold.

Our refoldability experiments utilize a frequent condition employed in classic protein folding studies, namely complete denaturation in 6 M guanidinium chloride, followed by rapid dilution to renaturing conditions. Clearly, these conditions do not mimic any normal physiological stressor and certainly not the slow progressive changes associated with aging. Nevertheless, we would argue that refoldability under these conditions is relevant because it reports on the ability of a protein to navigate to its native structure spontaneously and unassisted. Refoldable proteins can be viewed as potentially intrinsically more resistant to age-related structural changes, because no matter what types or magnitudes of stressors they may experience that temporarily result in unfolding or misfolding, they possess the inherent capacity to fold to their native structure autonomously. On the other hand, nonrefoldable proteins are more likely to be unable to recover from the damage accrued over time because they cannot efficiently self-correct them; hence, they are more likely to accumulate such deformations over age in the context of a disrupted proteostasis network. And while conventional thinking about the timescales associated with protein folding might assume that only covalent modifications would represent an accumulable form of damage over longer timescales, recent work on protein entanglement has demonstrated that certain misfolded conformations possessing changes in entanglement can be long-lived (*70*), and also correlate with nonrefoldability (*10*).

We note that the precise way proteins misfold upon attempted refolding from denaturant versus during aging are likely distinct in many cases. The association between CASC and nonrefoldability when looking at peptides specifically was not as robust as when considering entire proteins, as CASC sites are only somewhat more likely to be located near sites with altered structure after refolding. Though there are a few interesting exceptions (like DDX39B, cf. Fig. 3), this negative result is logical, as age-related protein misfolding would not be expected to occur from a globally denatured state, as it does in our *in vitro* refolding study. On the other hand, the data presented here present a case that nonrefoldable proteins, because of their incapacity to self-correct, are more likely to be permanently affected by stressors such as aging and hence provide a new window into the biophysics behind aging and proteostasis loss.

This study shows that soluble proteins with conformational differences exist between the hippocampus of rats with age-related spatial learning and memory impairment and those without, but at this stage, does not yet provide a mechanism for how these altered conformations form or persist. It is important to point out that CASC proteins are not amyloids. Due to their stability, amyloids are resistant to many degradation pathways (*71*), providing a rationale for why they can persist and accumulate over time during aging. However, using a bioinformatic predictor for amyloid formation propensity (*72*), we find that CASC proteins are not, in general, amyloidogenic (Fig. S8). Hence, an alternative model is required.

We put forward three potential explanations for how soluble misfolded conformations might populate and note at this stage that these are hypotheses that would need to be explored deeper with future study. These explanations are all related to proteostasis. One possibility is that CASC proteins are those which are more reliant on various molecular chaperones to properly fold. Rats with cognitive decline may have lower levels of certain chaperones or less functional chaperones, resulting in CASC proteins (which often contain complicated structural elements) to become the first “victims” to an impaired proteostasis network. Some chaperones have been found at lower levels in aged cells (*73*, *74*) and chaperone function have been linked to cognition (*75*).

A second possibility is that CASC proteins are those which are more dependent on cotranslational folding to fold. Recent studies have suggested that translation can be dysregulated in aged cells (*76*, *77*) and this could provide a global stress that is particularly acute for proteins sensitive to the precise timing of translation elongation or chaperones that act cotranslationally. Circumstantial evidence that supports this view is that CASC proteins tend to be nonrefoldable, which may be more reliant on cotranslational folding to bypass incorrect kinetically-trapped conformations (*38*). We also identified eukaryotic elongation factor 2 as a CASC. A structural change of such a critical protein for translation may have a functional effect on translation *in vivo*.

A third possibility is that CASC proteins represent conformations with impaired turnover, hence despite an altered non-native structure, they are degraded more slowly than their correctly-folded native forms, which results in a slow accumulation over the course of time. Protein clearance mechanisms, such as autophagy, are already known to be impaired with age (*15*), along with protein turnover in general (*78*). In support of this hypothesis, when we cross-compare our results to their mouse orthologs in a proteome-wide assessment of protein lifetimes in the mouse brain (*79*), we find a strong association wherein long lifetime proteins have a high propensity to be CASC (e.g., proteins with lifetimes >32 days are 30% CASC), and short lifetime proteins are less likely to be CASC (e.g., proteins with lifetimes <4 days are 7% CASC; Fig. S9). The trend is present when considering lifetimes of proteins from cortex homogenates or synaptic fractions, or proteins from old mice (aged 21 months) or young mice (aged 5 months; Fig. S9). This finding suggests that low lifetime proteins may be degraded before they have an opportunity to populate aberrant conformations, while proteins that persist for longer are at greater risk. Though protein turnover in the brain has been studied, previous experiments have not yet explored conformation-specific changes in turnover, which could prove to be an interesting area for future study. One hypothesis is that longer lifetime proteins might undergo misfolding events that further perturb their turnover, which would in turn provide a way for them to accumulate slowly over the course of aging. We caution however that the present study cannot yet establish whether the subpopulation of proteins that populate CASC conformations are older or newly synthesized.

A further caveat is that the subfield dissection approach used in the current study does not allow for definitive identification of where proteins are produced, only where they are localized. For example, the CA1 molecular layer would include axon terminals from CA3 Schaffer collaterals as well as from entorhinal cortex inputs. This leaves open the possibility that CASCs identified in the CA1 or DG preparations could have originated from upstream synaptic partners. If CASCs formed primarily at synapses, then the CASCs identified in the CA1 dissection could have originated from cell bodies of CA3 neurons. Previous studies identified greater changes in transcription and neuronal hyperactivity in CA3 in AI rats (*6*, *25*, *29*, *46*); further research will be needed to better elucidate if changes in protein structure in CA1 are connected to the other changes observed in CA3 neurons.

To summarize, we have found that structural proteomics is an emerging “omics” method that is particularly relevant for dissecting the features associated with progressive age-related cognition loss, providing molecular details to the sporadic changes that bring on cognitive decline. Modern methods such as LiP-MS are appropriately sensitive to detect subtle structural alterations in soluble misfolded proteins, which are pervasive in aged rats’ hippocampi. Mechanistic and structural studies on specific CASC proteins could promote some of the proteins disclosed here to be considered as therapeutic targets or early disease biomarkers.

## MATERIALS AND METHODS

### 1. Animal Care and Tissue Collection

Long-Evans rats were obtained from Charles River Laboratories. Aged (AU and AI) rats were obtained as retired breeders at approximately age 6-9 months and housed in a vivarium at Johns Hopkins University until 24 months of age. Young rats were also obtained from Charles River Laboratories at approximately age 4-6 months and housed in the same vivarium. Food and water were given ad libitum and animals were on a 12-hour light/dark cycle. Animals housing and use was performed in accordance with protocols approved by the Johns Hopkins University Institutional Animal Care and Use Committee.

Twenty animals were used for the CA1 CASC LiP experiment (10 AU, 7 AI, and 3 Y). The DG and CA3 studies used the same animals, except omitting one AU animal for a total of 9 AU, 7 AI, and 3 Y. Two weeks after completion of behavioral testing, all rats were deeply anesthetized with isoflurane and sacrificed by rapid decapitation. The two-week time frame was chosen to maximize the likelihood of detecting stable, baseline protein profile differences and minimizing the effects of learning or swimming induced differences. Brains were immediately extracted and placed in ice-cold phosphate-buffered saline (PBS). The hippocampus was separated from the brain and, for the CASC LiP study, the CA1, CA3, and DG were microdissected by hand from 400 micron transverse sections of the hippocampus along its entire longitudinal extent. Hippocampi for the refoldability study were not subfield dissected (*80*). Tissue was snap frozen on dry ice and moved to a -80°C freezer for storage.

### 2. Morris Water Maze Cognitive Assessment

Behavioral assessment of memory function in a Morris water maze task was conducted as previously described (*31*). Briefly, rats were trained for eight days (three trials per day) to locate a camouflaged escape platform that remained at the same location throughout training in a water maze. Every sixth trial consisted of a probe trial (no escape platform for the first 30s of the trial) that served to assess the development of a spatially localized search. Learning Index scores were derived from each rat’s proximity to the platform during probe trials, with lower scores reflecting a more accurate search indicating better retention of the platform location. A learning index cutoff of 240 was used to identify aged rats as Aged Unimpaired (AU) or Aged Impaired (AI), with higher scores representing worse performance and reflecting scores that fall inside or outside the normative range collected from young adult Long Evans rats over many years. Cue training was used to assess the sensorimotor and motivational status of the rats. Only rats with successful cue training performance were included in the present study.

### 3. Aging/Cognition Structural Proteomics Experiment 3a. Limited Proteolysis (LiP)

Frozen hippocampal subfields were removed from the -80°C freezer and immediately added to 1mL of chilled lysis buffer (20mM Tris pH 8, 150mM KCl, 10mM NaCl, 2mM MgCl_2_) containing protease inhibitors (phenylmethylsulfonyl fluoride (PMSF) 0.5mM final concentration (f.c.), bestatin 0.05mM f.c., E64 0.015mM f.c., all diluted from 100X stock in DMSO) and DNase (0.1 mg/mL f.c., diluted from 100X stock in Millipore water (MPW)) in a 1mL dounce homogenizer (Wheaton USA) on ice. Samples were vigorously homogenized on ice. Homogenized tissue was moved to a 1.5mL microfuge tube and clarified at 15000xg for 15 min at 4°C to remove insoluble cellular debris. The supernatant was moved to a fresh 1.5mL microfuge tube and allowed to sit at room temp for at least two hours and twenty minutes before limited proteolysis (LiP), to ensure that PMSF had sufficient time to hydrolyze (approx. four half lives) and would not inhibit our protease during the LiP step. During this time, protein concentrations of the samples were determined using the bicinchoninic acid (BCA) assay, according to the manufacturer’s protocol (Pierce Rapid Gold Protein Assay Kit, Thermo Scientific A53225) and using bovine serum albumin (BSA) as a standard. Lysates were normalized to 1 mg/mL or 0.5 mg/mL by dilution with lysis buffer.

After adequate time for hydrolysis of PMSF has passed, LiP can be performed. Proteinase K (PK, Thermo Scientific 17916, previously prepared as 1 mg/mL stock in 1:1 v/v of lysis buffer to 20% glycerol, aliquoted, flash frozen and stored at -20°C) was diluted such that 2uL of the dilution would contain 1:100 w/w of PK to the amount of sample to be digested (in this case, either 1μg or 2μg for 100μg or 200μg sample, respectively). 2μL of appropriately diluted PK was placed at the bottom of a fresh 1.5mL microfuge tube. 200μL of normalized sample was added to the tube containing 2μL PK and pipetted rapidly up and down seven times to mix thoroughly. This mixture was incubated for exactly one minute before the tube was added to a 105°C mineral oil bath for five minutes to quench the PK reaction. For each sample, a corresponding no-PK, non-LiP control was performed. In this case, 200μL of normalized lysate was added to a fresh 1.5mL tube without PK and then added to the oil bath for 5 minutes. After 5 minutes, all samples were removed from the oil bath and quickly centrifuged to collect condensation and transferred to a new 2mL tube containing 152mg urea (8M urea f.c., 314μL f.v.).

### 3b. Mass Spectrometry Sample Preparation

To each sample in 8M urea, in order to reduce disulfides, dithiothreitol (DTT) was added to 10mM f.c. (4.5μL added from a freshly made 700mM DTT stock) and incubated in a thermomixer for 30 min at 37°C and 700 rpm. To cap cysteines, iodoacetamide (IAA) was added to a f.c. of 40mM (18μL added from a freshly made 700mM IAA stock) and incubated at room temperature (rt) in the dark for 45 min. Samples were diluted to 2M urea f.c. by adding 1010μL of a freshly-made 100mM ammonium bicarbonate (ambic) stock. To each sample, 2μg trypsin (New England Biolabs) was added (1:50 w/w of trypsin to protein). Samples were incubated overnight (approximately 16 hours) in a thermomixer at 25°C and 700 rpm. Digested samples were acidified with trifluoroacetic acid (TFA, Acros) to a 1% f.c. by volume by adding 16.6μL TFA. Peptides were desalted using Sep-Pak Vac 1cc (50mg) C18 cartridges (Waters). Cartridges were placed on a vacuum manifold and conditioned by adding 2×1mL Buffer B (0.5% TFA in 80% liquid chromatography (LC) grade acetonitrile (ACN) and 20% LC grade water). Cartridges were equilibrated with 4×1mL Buffer A (0.5% TFA in LC grade water). Peptide samples were loaded on each cartridge. Cartridges were washed with 4×1mL Buffer A. Cartridges were removed from the vacuum manifold, placed in a 15mL conical tube, and eluted by adding 1mL Buffer B to the cartridge and spinning in a swing bucket rotor (Eppendorf 5910 R) at 300rpm for 5 min. Eluent was transferred from the conical vial to a fresh 1.5mL microfuge tube and dried using a vacuum centrifuge (Eppendorf Vacufuge Plus). Dried peptides were stored at -80°C until ready for analysis.

### 3c. Mass Spectrometry Data Acquisition

Peptide samples were resuspended in 0.1% formic acid (Optima, Fisher) in LC grade water (Optima, Fisher) to a f.c. of 1μg/μL. Approximately 1μg of peptides were injected into a Thermo UltiMate 3000 UHPLC system for chromatographic separation. The column temperature was maintained at 40 °C, and the flow rate was 0.300μL/min for the duration of the run. Solvent A (0.1% formic acid (FA)) and Solvent B (0.1% FA in ACN) were used as the chromatography solvents. Peptides were allowed to accumulate onto the trap column (Acclaim PepMap 100, C18, 75μm x 2cm, 3μm, 100Å column) for 10 min (during which the column was held at 2% Solvent B). The peptides were resolved by switching the trap column to be in-line with the separating column (Acclaim PepMap RSLC, C18, 75μm x 2cm, 2μm, 100Å column), increasing the gradient to 5% B over 5 min and then applying a 95 min linear gradient from 5% B to 25% B. Then the gradient was increased linearly to 40% B over 25 minutes. Next, the gradient was held at 40% B for 5 min and then increased again from 40% B to 90% B. The gradient was held at 90% B for 5 min. The column was then cleaned with a sawtooth gradient to purge residual peptides between runs in a sequence. A one-hour blank cleaning sequence was also run between every 3-6 samples to ensure a clean column. A Thermo Q Exactive HF-X Orbitrap mass spectrometer was used to analyze protein digests. A full MS scan in positive ion mode was followed by 20 data-dependent MS scans. The full MS scan was collected using a resolution of 120000 (@ m/z 200), an AGC target of 3E6, a maximum injection time of 64ms, and a scan range from 350 to 1500 m/z. The data-dependent scans were collected with a resolution of 15000 (@ m/z 200), an AGC target of 1E5, a minimum AGC target of 8E3, a maximum ion injection time of 55ms, and an isolation window of 1.4 m/z units. To dissociate precursors prior to their reanalysis by MS2, peptides were subjected to an HCD of 27% normalized collision energies. Fragments with charges of 1, 6 or higher, and unassigned were excluded from analysis, and a dynamic exclusion window of 30.0s was used for the data-dependent scans.

### 3d. Data Analysis of Individual Runs

For batch analyses of individual runs, Proteome Discoverer (PD, version 2.4, Thermo Fisher) was used. These analyses were performed to gather quality metrics such as number of Peptide Spectrum Matches (PSMs) and percent half-trypticity.

Default settings were used except where specified otherwise. The data were searched against the FASTA reference proteome for rats, *Rattus norvegicus* (UP000002494, Uniprot.org). The Sequest HT node was used for peptide identification, and the Percolator node was used for false discovery rate (FDR) validation. For Sequest HT, a semi-specific trypsin search was performed, allowing for 2 missed cleavages. Peptides searched for had a minimum length of 6 and maximum length of 144 amino acids. A precursor mass tolerance of 10ppm for the MS1 level and a fragment mass tolerance of 0.02Da for the MS2 level. Carbidomethylation of cysteines was set as a static modification and oxidation of methionine was set as a dynamic modification. For Percolator, FDR targets were set to 0.01 (strict) and 0.05 (relaxed). Data were exported from the .pdResult file to excel in a two-tier file, containing proteins with their respective peptide groups underneath them. These data were analyzed using an in-house Python script. Briefly, the program counts, for each PD output file, the number of proteins and peptides, and then determines whether each peptide is tryptic, half-tryptic, or ambiguous. Half-tryptic peptides are peptides with one end immediately preceding arginine or lysine (the two amino acids after which trypsin cuts). Fully tryptic peptides have both ends immediately preceded by arginine or lysine. Ambiguous peptides occur when the amino acid immediately prior to or immediately after the peptide of interest could be multiple possible amino acids (as indicated by PD). In some cases, it is not possible to determine if the peptide is fully tryptic or half-tryptic.

### 3e. Label Free Quantification and LiP-MS Data Analysis

Two label-free quantification (LFQ) analyses were performed, one with all the no-PK (trypsin only) controls, and one with the LiP samples (which contain PK). MSFragger and the Minora Feature Detector nodes in PD were used for the LFQs. Default settings were used except where specified otherwise. The data were searched against the FASTA reference proteome for rats, *Rattus norvegicus* (UP000002494, Uniprot.org). In the MSFragger node for peptide identification, semi-specific trypsin searches were performed, with up to 2 missed cleavages allowed. Fragment mass tolerance was 20ppm. Peptide masses were set to a minimum of 500Da and a maximum of 5000Da, and peptides of a minimum length of 7 amino acids and a maximum length of 50 amino acids were allowed. Carbidomethylation of cysteine was set as a static modification, and methionine oxidation and N-terminal acetylation were set as dynamic modifications. The maximum charge state for theoretical fragments was 2. The Philosopher node was used for FDR validation.

One of the LFQs, the control LFQ, compared all non-PK, trypsin-only, samples. The LiP LFQ compared all LiP samples, where PK was added. For each LFQ, samples were categorized as either aged unimpaired (AU), aged impaired (AI), or young (Y), and ratios were generated for AU/AI, AI/Y, and AU/Y. For the control LFQ, raw normalized extracted ion intensity data for the identified peptides were exported from the .pdResult file to a three-tier excel file (protein with peptide groups under each protein, with consensus features underneath each peptide group). For the LiP LFQ, ion intensity data was exported in a two-tier excel file, with peptide groups exported with consensus features underneath each peptide group. Also for the LiP LFQ, another two-tier file with peptide groups and the protein(s) they match to underneath each peptide group was exported. This two two-tier export approach was used because with three-tier files, peptides that matched to multiple proteins would be repeated underneath the other proteins it matched to.

These data were further processed using an in-house python script. Briefly, normalized ion counts were collected across the AU, AI, and Y replicates for each peptide group. Effect sizes are the ratio of averages (reported in log2), and P-values (reported as –log10) were assessed using t tests with Welch’s correction for unequal population variances. Missing data are treated in a special manner. If a feature is not detected in only one injection of that category, the missing value is dropped. If a feature is not detected in more than one injection of any one category (ie AU, AI, or Y), we use those data, and fill the missing values with 1000 (the ion limit of detection for this mass analyzer). In many situations, our data provide multiple independent sets of quantifications for the same peptide group. This happens most frequently because the peptide is detected in multiple charge states. In this case, we calculate effect size and P-value for all features that map to the same peptide group. If the features all agree with each other in sign, they are combined: the quantification associated with the median among available features is used and the P-values are combined with Fisher’s method. If the features disagree with each other in sign, the P-value is set to 1. Coefficients of variation (CV) for the peptide abundance in the replicates within each category are also calculated. For each peptide, the protein ratio from the control LFQ is referenced – if the protein abundance ratio is greater than two-fold in either direction, the P-value for the protein ratio is less than 0.01, and the peptide matches to a single protein, the peptide ratio is normalized by the protein abundance ratio. The script returns a file listing all the peptides that can be confidently quantified, and provides the effect sizes and P-values for all three ratios (AU/AI, AI/Y, and AU/Y), CVs for each category, proteinase K site (if half-tryptic) for the peptide, and associated protein metadata.

### 3f. CASC/Hit Determination

From the list of peptides and AU/AI, AI/Y, and AU/Y ratios generated by the program, we generate a list of proteins. For each protein, the number of peptides that match to that protein for each ratio that have nonzero normalized peptide ratios and P-values is considered the number of total peptides. Peptides are considered significantly different for the ratio of interest if the normalized peptide ratio effect size is 2 or greater (the peptide is twice or more abundant in one of the two categories in the ratio compared to the other) and the P-value is less than 0.01. If the effect size is greater than 64, a P-value of less than 0.0158 is acceptable to be considered significant. If a peptide can be matched to only a single protein, the peptide is considered “unique”. If the peptide matches to multiple proteins, it is not considered unique. The “unique” tally counts all unique peptides for that protein in that ratio, and all unique significant peptides for that ratio. The “all” category is a total of both non-unique peptides that match to multiple proteins, and “unique” peptides. The “all” tally counts all peptides for that protein in that ratio, and all significant peptides for that ratio. A protein is considered to be a hit in our study, meaning there is likely a structural difference between the two categories in the ratio, if there is more than one peptide detected that matches to said protein, and of those peptides, more than one is significantly different. A Cognition-Associated Structural Change (CASC) is a protein that is considered a hit for the AU/AI ratio.

### 4. Aggregation Assay

#### 4a. Preparation of Native and Refolded Hippocampal Extracts

Frozen whole hippocampi from young (approx. 6 month old) rats were removed from the -80°C freezer and immediately added to 750μL of chilled lysis buffer (20mM Tris pH 8, 100mM NaCl, 2mM MgCl_2_) containing protease inhibitors (PMSF 0.5mM final concentration (f.c.), bestatin 0.05mM f.c., E64 0.015mM f.c., all diluted from 100X stock in DMSO) and DNase (0.1 mg/mL f.c., diluted from 100X stock in MPW) in a 1mL dounce homogenizer (Wheaton USA) on ice. Samples were vigorously homogenized on ice. Homogenized tissue was moved to a 1.5mL microfuge tube and clarified at 16000xg for 15 min at 4°C to remove insoluble cellular debris. The supernatant was moved to a fresh 1.5mL microfuge tube and kept on ice. Protein concentrations of the samples were determined using the bicinchoninic acid (BCA) assay, according to the manufacturer’s protocol (Pierce Rapid Gold Protein Assay Kit, Thermo Scientific A53225) and using bovine serum albumin (BSA) as a standard. Lysates were normalized to 2 mg/mL by dilution with lysis buffer.

To prepare native samples, normalized lysate was diluted to 0.115 mg/mL f.c. using native dilution buffer (20mM Tris pH 8, 100mM NaCl, 2mM MgCl_2_, 1.061mM DTT, 0.0637M guanidinium chloride (GdmCl)). This buffer has a small amount of GdmCl to match the concentration of GdmCl that will be present after refolding of unfolded samples. Final concentrations of all components were 0.115 mg/mL protein, 20mM Tris pH 8, 100mM NaCl, 2mM MgCl_2_, 0.06mM GdmCl, and 1mM DTT. Native samples were incubated at room temp overnight.

To prepare refolding samples, proteins were first unfolded. Initially, 500μg of protein was added to a fresh 1.5mL microfuge tube, and 25mg of GdmCl powder and 0.61μL of a 700mM DTT stock was added. Refolding samples were placed in a Vacufuge Plus (Eppendorf) and volume was reduced to 43μL. Final concentrations of all components were 11.5 mg/mL protein, 115mM Tris pH 8, 575mM NaCl, 11.5mM MgCl_2_, 6M GdmCl, and 10mM DTT. Samples were incubated at room temp overnight to ensure complete unfolding of the samples.

To refold the samples, samples were diluted 100X in refolding dilution buffer (19.03mM Tris pH 8, 95.14mM NaCl, 1.9mM MgCl_2_, 0.909mM DTT) by adding 10μL of unfolded sample into 990μL of refolding dilution buffer. Final concentrations of all components were 0.115 mg/mL protein, 20mM Tris pH 8, 100mM NaCl, 2mM MgCl_2_, 0.06mM GdmCl, and 1mM DTT. All concentrations are the same as the native samples. Refolded samples were incubated for two hours at room temp to allow proteins to refold.

#### 4b. Determination of Aggregation Percentage

Native and refolded samples were centrifuged at 16000xg for 15 min at 4°C to collect any aggregated material. The supernatant was removed, taking care not to disturb the pellet containing aggregated proteins. The pellet was washed with 200μL lysis buffer, which reduces the potential interference from reducing agents in the sample with the BCA assay. The pellet was resuspended in 75μL of 8M urea in MPW. Protein concentrations were determined by the BCA assay, as previously described. There were three biological replicates for the native and refolded samples, and the BCA was performed with technical triplicates. The percent aggregation was determined by determining the total amount of protein in the pellet and dividing by the total amount of protein in the initial native or refolded sample. This calculation yields the percent protein that was precipitated in the sample. Data are reported as mean ± standard deviation. T-tests were performed in Prism 10 (Graphpad) using a parametric unpaired T-test with Welch’s correction for unequal population variances (the same standard deviation is not assumed).

### 5. Refoldability Assay

#### 5a. Preparation of Native and Refolded Hippocampal Extracts

Three frozen whole hippocampi from young (approx. 6 month old) rats were removed from the -80°C freezer and each immediately added to 1mL of chilled lysis buffer (20mM Tris pH 8, 100mM NaCl, 2mM MgCl_2_) containing protease inhibitors (PMSF 0.5mM final concentration (f.c.), bestatin 0.05mM f.c., E64 0.015mM f.c., all diluted from 100X stock in DMSO) and DNase (0.1 mg/mL f.c., diluted from 100X stock in MPW) in a 1mL dounce homogenizer (Wheaton USA) on ice. Samples were vigorously homogenized on ice. Homogenized tissue was moved to a 1.5mL microfuge tube and clarified at 16000xg for 15 min at 4°C to remove insoluble cellular debris. The supernatant was moved to a fresh 1.5mL microfuge tube and kept on ice. Protein concentrations of the samples were determined using the BCA assay, according to the manufacturer’s protocol (Pierce Rapid Gold Protein Assay Kit, Thermo Scientific A53225) and using BSA as a standard. Lysates were normalized to 2 mg/mL by dilution with lysis buffer.

To prepare native samples, normalized lysate was diluted to 0.115 mg/mL f.c. using native dilution buffer (20mM Tris pH 8, 100mM NaCl, 2mM MgCl_2_, 1.061mM DTT, 0.0637M guanidinium chloride (GdmCl)) by adding 11.5μL of 2 mg/mL lysate to 188.5μL of native dilution buffer. This buffer has a small amount of GdmCl to match the concentration of GdmCl that will be present after refolding of unfolded samples. Final concentrations of all components were 0.115 mg/mL protein, 20mM Tris pH 8, 100mM NaCl, 2mM MgCl_2_, 0.06mM GdmCl, and 1mM DTT. Native samples were incubated at room temp for one hour before proceeding to limited proteolysis (LiP). Importantly, 2.5 hours must have passed since initial addition of PMSF in order to ensure it is hydrolyzed sufficiently to not interfere with PK in the LiP step.

To prepare refolding samples, proteins were first unfolded. Initially, 500μg of protein (250 μL of 2 mg/mL lysate) was added to a fresh 1.5mL microfuge tube, and 25mg of GdmCl powder and 0.61μL of a 700mM DTT stock was added. Refolding samples were placed in a Vacufuge Plus (Eppendorf) and volume was reduced to 43μL. Final concentrations of all components were 11.5 mg/mL protein, 115mM Tris pH 8, 575mM NaCl, 11.5mM MgCl_2_, 6M GdmCl, and 10mM DTT. Samples were incubated at room temp overnight to ensure complete unfolding of the samples.

To refold the samples, samples were diluted 100X in refolding dilution buffer (19.03mM Tris pH 8, 95.14mM NaCl, 1.9mM MgCl_2_, 0.909mM DTT) by adding 10μL of unfolded sample into 990μL of refolding dilution buffer. Final concentrations of all components were 0.115 mg/mL protein, 20mM Tris pH 8, 100mM NaCl, 2mM MgCl_2_, 0.06mM GdmCl, and 1mM DTT. All concentrations are the same as the native samples. Refolded samples were incubated for either 1 min, 5 min, or 2 hours at room temp to allow proteins to refold before proceeding to the LiP step.

#### 5b. Limited Proteolysis

After the prescribed time had passed (either 1 min, 5 min, or 2 hours from refolding for refolded samples, and 1hr from native dilution for native samples), LiP was performed. Proteinase K (PK, Thermo Scientific 17916, previously prepared as 1 mg/mL stock in 1:1 v/v of lysis buffer to 20% glycerol, aliquoted, flash frozen and stored at - 20°C) was diluted such that 2uL of the dilution would contain 1:100 w/w of PK to the amount of sample to be digested (in this case, 0.23μg for 23μg of sample). 2μL of appropriately diluted PK (0.115 mg/mL) was placed at the bottom of a fresh 1.5mL microfuge tube. 200μL of sample was added to the tube containing 2μL PK and mixed by a quick vortex followed by a short spin in a benchtop centrifuge to ensure all sample is at the bottom of the tube. This mixture was incubated for exactly one minute (including vortex and centrifugation time) before the tube was added to a 105°C mineral oil bath for five minutes to quench the PK reaction. For each native sample and 2hr refolding sample, a corresponding no-PK, non-LiP control was performed (6 total). In this case, 200μL of normalized lysate was added to a fresh 1.5mL tube without PK and then added to the oil bath for 5 minutes. After 5 minutes, all samples were removed from the oil bath and quickly centrifuged to collect condensation and transferred to a new 2mL tube containing 152mg urea (8M urea f.c., 314μL f.v.).

#### 5c. Mass Spectrometry Sample Preparation

To each sample in 8M urea, in order to reduce disulfides, dithiothreitol (DTT) was added to 10mM f.c. (4.5μL added from a freshly made 700mM DTT stock) and incubated in a thermomixer for 30 min at 37°C and 700 rpm. To cap cysteines, iodoacetamide (IAA) was added to a f.c. of 40mM (18μL added from a freshly made 700mM IAA stock) and incubated at room temperature (rt) in the dark for 45 min.

Samples were diluted to 2M urea f.c. by adding 942μL of a freshly-made 100mM ammonium bicarbonate (ambic) stock. To each sample, 0.46μg trypsin (New England Biolabs) was added (1:50 w/w of trypsin to protein). Samples were incubated overnight (approximately 16 hours) in a thermomixer at 25°C and 700 rpm. Digested samples were acidified with trifluoroacetic acid (TFA, Acros) to a 1% f.c. by volume by adding 12.8μL TFA. Peptides were desalted using Sep-Pak Vac 1cc (50mg) C18 cartridges (Waters). Cartridges were placed on a vacuum manifold and conditioned by adding 2×1mL Buffer B (0.5% TFA in 80% liquid chromatography (LC) grade acetonitrile (ACN) and 20% LC grade water). Cartridges were equilibrated with 4×1mL Buffer A (0.5% TFA in LC grade water). Peptide samples were loaded on each cartridge. Cartridges were washed with 4×1mL Buffer A. Cartridges were removed from the vacuum manifold, placed in a 15mL conical tube, and eluted by adding 1mL Buffer B to the cartridge and spinning in a swing bucket rotor (Eppendorf 5910 R) at 300rpm for 5 min. Eluent was transferred from the conical vial to a fresh 1.5mL microfuge tube and dried using a vacuum centrifuge (Eppendorf Vacufuge Plus). Dried peptides were stored at -80°C until ready for analysis.

#### 5d. Mass Spectrometry Data Acquisition

Data acquisition was the same as reported in section 3c, except that peptides were resuspended in 0.1% formic acid (Optima, Fisher) in LC grade water (Optima, Fisher) to a f.c. of 0.5μg/μL (instead of 1μg/μL), so a higher injection volume was used to still inject approximately 1μg of peptides.

#### 5e. Data Analysis of Individual Runs

For batch analyses of individual runs, analyses were performed the same as in section 3d.

#### 5f. Label Free Q3uantification and LiP-MS Data Analysis

Four label-free quantification (LFQ) analyses were performed, one with all the no-PK (trypsin only) controls, and three with the LiP samples (which contain PK), one for each refolding timepoint. MSFragger and the Minora Feature Detector nodes in PD were used for the LFQs. Default settings were used except where specified otherwise. The data were searched against the FASTA reference proteome for rats, *Rattus norvegicus* (UP000002494, Uniprot.org). In the MSFragger node for peptide identification, semi-specific trypsin searches were performed, with up to 2 missed cleavages allowed. Fragment mass tolerance was 20ppm. Peptide masses were set to a minimum of 500Da and a maximum of 5000Da, and peptides of a minimum length of 7 amino acids and a maximum length of 50 amino acids were allowed. Carbidomethylation of cysteine was set as a static modification, and methionine oxidation and N-terminal acetylation were set as dynamic modifications. The maximum charge state for theoretical fragments was 2. The Philosopher node was used for FDR validation.

The control LFQ compared all non-PK, trypsin-only samples, three from the native control and three from the 2hr refolding controls. The LiP LFQs compared all LiP samples, where PK was added, one for each timepoint, where each compared the native LiP samples and the refolded samples for that timepoint. For each LFQ, samples were categorized as either native or refolded, and a ratio was generated for refolded/native. For the control LFQ, raw normalized extracted ion intensity data for the identified peptides were exported from the .pdResult file to a three-tier excel file (protein with peptide groups under each protein, with consensus features underneath each peptide group). For the LiP LFQ, ion intensity data was exported in a two-tier excel file, with peptide groups exported with consensus features underneath each peptide group. Also for the LiP LFQ, another two-tier file with peptide groups and the protein(s) they match to underneath each peptide group was exported. This two two-tier export approach was used because with three-tier files, peptides that matched to multiple proteins would be repeated underneath the other proteins it matched to.

These data were further processed using an in-house python script. Briefly, normalized ion counts were collected across the native and refolded replicates for each peptide group. Effect sizes are the ratio of averages (reported in log2), and P-values (reported as –log10) were assessed using t tests with Welch’s correction for unequal population variances. Missing data are treated in a special manner. If a feature is not detected in only one injection of all six (both native and refolded), the missing value is dropped. If a feature is not detected in all three native or all three refolded samples, but is detected in all three of the other type of samples (native or refolded), the three missing values are replaced with a value of 1000 (the ion limit of detection). Any other permutation of missing data (such as missing in two injections) results in the disregard of that quantification. In many situations, our data provide multiple independent sets of quantifications for the same peptide group. This happens most frequently because the peptide is detected in multiple charge states. In this case, we calculate effect size and P-value for all features that map to the same peptide group. If the features all agree with each other in sign, they are combined: the quantification associated with the median among available features is used and the P-values are combined with Fisher’s method. If the features disagree with each other in sign, the P-value is set to 1. Coefficients of variation (CV) for the peptide abundance in the replicates within each category are also calculated. For each peptide, the protein ratio from the control LFQ is referenced – if the protein abundance ratio is greater than two-fold in either direction, the P-value for the protein ratio is less than 0.01, and the peptide matches to a single protein, the peptide ratio is normalized by the protein abundance ratio. The script returns a file listing all the peptides that can be confidently quantified, and provides the effect sizes and P-values for the refolded/native ratio, the CV, proteinase K site (if half-tryptic) for the peptide, and associated protein metadata.

#### 5g. Refoldable/Nonrefoldable Classification

From the list of peptides and the refolded/native ratio generated by the program, we generate a list of proteins. For each protein, the number of peptides that match to that protein that have a nonzero normalized peptide ratio and P-value is considered the number of total peptides. Peptides are considered significantly different if the normalized peptide ratio effect size is 2 or greater (the peptide is twice or more abundant in either the native or refolded samples) and the P-value is less than 0.01. If the effect size is greater than 64, a P-value of less than 0.0158 is acceptable to be considered significant. If a peptide can be matched to only a single protein, the peptide is considered “unique”. If the peptide matches to multiple proteins, it is not considered unique. The “unique” tally counts all unique peptides for that protein and all unique significant peptides. The “all” category is a total of both non-unique peptides that match to multiple proteins and “unique” peptides. The “all” tally counts all peptides for that protein and all significant peptides. A protein is considered to be nonrefoldable, meaning there is likely a structural difference between the refolded protein and the native protein, if there is more than one peptide detected that matches to said protein, and of those peptides, more than one is significantly different. A refoldable protein, where the native and refolded protein likely have the same structure, has more than one peptide detected that matches to the protein, but no more than one significantly different peptide. Proteins with only one peptide matched are not considered to have enough information to be categorized as either refoldable or nonrefoldable.

### 6. PTM Analysis

#### 6a. Re-Searches of Data for PTMs

In order to determine if PTMs confounded the LiP data analysis, we first re-searched all control (no PK added) samples for each region to look for PTMs, specifically phosphorylation and oxidation modifications. Proteome Discoverer (PD, version 2.4, Thermo Fisher) was used with default settings, except where specified otherwise. The data were searched against the FASTA reference proteome for rats, *Rattus norvegicus* (UP000002494, Uniprot.org). The Sequest HT node was used for peptide identification, and the Percolator node was used for false discovery rate (FDR) validation. For Sequest HT, settings were the same as in section 3d, except for modifications. We searched for phosphorylation and oxidation modifications in separate searches. For the phosphorylation analysis, we searched for dynamic phospho (+79.966 Da) modifications on S, T, and Y residues. We also allowed methionine oxidation (+15.995 Da) as a dynamic modification. For the oxidation analysis, we allowed for dynamic oxidation modifications (+15.995 Da) on methionine, tryptophan, phenylalanine, and tyrosine. For both searches, cysteine carbidomethylation was set as a static modification. For Percolator, FDR targets were set to 0.01 (strict) and 0.05 (relaxed). Peptide data were exported from the .pdResult file to excel. For oxidation, a two-tier file of peptides, then PSMs, was exported, as well.

Since oxidation is known to occur as a result of electrospray ionization (ESI) (*81*), a retention time correction was applied to oxidation analyses, to exclude oxidized peptides that are most likely artifacts of ionization. Peptides that are oxidized *in vivo* will have different retention times than non-oxidized peptides of the same sequence, whereas peptides oxidized during ESI, which occurs after LC separation, should have the same retention time as non-oxidized peptides of the same sequence. If an oxidized peptide’s average retention time was within ±1 min of the retention time of the non-oxidized peptide of the same sequence, that oxidation was considered to be an artifact and removed.

Percent phosphorylated and percent oxidized peptides were determined by dividing the number of peptides with one (or more) of the modification by the total number of peptides detected.

## 6b. Proximity Analysis

The number of CASC and non-CASC sites near a modification (either phosphorylation or oxidation, regarded separately) was determined using an in-house python script. All sites from the CASC study were considered, meaning all cut sites from proteinase K and all centers of fully tryptic peptides. Peptides are considered significantly different (CASC) if the normalized peptide ratio effect size is 2 or greater between AU and AI, and the P-value is less than 0.01. If the effect size is greater than 64, a P-value of less than 0.0158 is acceptable to be considered significant. Otherwise, they are considered non-CASC sites. Next, for each significant site from the CASC study, we asked if it was found within 15 amino acids on either side of a modification from the re-searched data, or if it was not. We repeated this with all non-significant sites from the CASC study. This was performed separately for each of the three hippocampal regions.

### 7. Bioinformatics

#### 7a. Collection of Protein Metadata

Molecular weight and isoelectric point of proteins were calculated by Proteome Discoverer (version 2.4, Thermo Fisher). Percent disorder was calculated by Metapredict (*39*). Domains for each protein were assigned using DomainMapper (*82*), which uses domain definitions from Evolutionary Classifications of Domains (ECOD) (*40*). Amyloidogenicity was assessed via PLAAC, a tool which looks for prion-like amino acid composition (*72*). Comparisons to turnover used protein turnover data from Kluever *et al.* (*79*).

#### 7b. GO Analyses

All gene ontology (GO) analyses were performed using ShinyGO version 0.77 (*56*). The species was set to rat (Rattus norvegicus, assembly name “Rat genes mRatBN7.2”, taxonomy ID 10116, sourced from ENSEMBL (*83*)). Settings for all searches were FDR cutoff of 0.05, minimum pathway size of 2, maximum pathway size of 2000, and redundancy removed. For the analysis, a list of accession numbers for proteins with more than one significant peptide (for either the AU/AI comparison or the 5 min refoldability native to refolded comparison) was supplied to ShinyGO, along with a reference set of the accession numbers of all proteins from the appropriate data set with greater than one peptide detected. We examined biological process, molecular function, and cellular component GO terms, in addition to Kyoto Encyclopedia of Genes and Genomes (KEGG) pathways (*66*), and disease pathways from rat genome database (RGD) (*84*).

#### 7c. CASC/Refoldability Overlap and Proximity Analysis

CA1 CASC and 5 minute refoldability overlap was determined by considering all proteins which are considered CASC (or nonrefoldable, for the refoldability study). A protein is considered CASC or nonrefoldable if there is more than one peptide for that protein where the normalized peptide ratio effect size is 2 or greater (between AU and AI for CASC, between 5 min refolded and native for refoldability), and the P-value is less than 0.01. If the effect size is greater than 64, a P-value of less than 0.0158 is acceptable to be considered significant. Other proteins are considered non-CASC (or refoldable) if they have greater than 1 peptide, but one or fewer are significantly different. The overlap was considered to be proteins in the intersection of sets of the appropriate category (for example, the intersection of the set of CA1 CASC proteins and the set of 5 min nonrefoldable proteins). Proteins that were only identified in either the CA1 CASC study or the 5 min refoldability study, but not in the other, were not considered in this analysis.

For the CASC and refold proximity analysis, we considered sites from the CA1 CASC study and the 5 minute refoldability study. We considered sites from the CA1 CASC study to be the location of the PK cut site of the significant peptide, or the center of the significant peptide if it is fully tryptic. For each significant site, we asked if the site is located within 5 amino acids on either side of a nonrefoldable site, then if is within 5 amino acids on either side of a refoldable site, or if it is not located near either. We then asked this for nonsignificant sites, as well. Nonrefoldable and refoldable sites were considered to be significant and non-significant peptides, respectively, from the 5 min refoldability study.

#### 7d. Structural Visualization

Protein structures were generated from AlphaFold version 2 (*43*) from their accession numbers. Accession numbers used were P62260 (14-3-3 epsilon), Q63413 (DDX39B), P61765 (syntaxin-binding protein 1), P05197 (eukaryotic elongation factor 2), and P50137 (transketolase 1). Structures were visualized using PyMOL version 1.8.3.2. Significant sites from the CA1 CASC study (PK cut sites of significant peptides, or centers of fully tryptic significant peptides) were represented as light blue spheres on the structure. Significant sites from the 5 min refoldability study (nonrefoldable sites) were represented as black spheres on the structure. Sites that were significant in both the CA1 CASC study and 5 min refoldability study were represented as dark blue spheres. Sequence diagrams were also generated, with the domains identified by DomainMapper (*82*) represented. The significant sites were shown as vertical lines of the same colors as the corresponding spheres (light blue for significant CA1 CASC site, black for significant 5 min nonrefoldable sites, and dark blue for sites that are significant in both studies).

### 8. Reproducibility Experiment

Three AU hippocampi with subfield dissections were obtained in order to determine biological variation (between the three samples) and technical variation (of LiP, performed multiple times on the same lysate, and of the instrument, by shooting identical samples more than once). Each CA1 region was lysed by Dounce homogenization vigorously on ice in 0.5mL chilled lysis buffer (20mM Tris pH 8.0, 100mM NaCl, 2mM MgCl_2_) in the presence of protease inhibitors (0.5mM PMSF, 0.015mM E64, 0.05mM bestatin f.c.) and 0.1 mg/mL f.c. DNase. Lysate was clarified for 15 min at 15000xg and 4°C. remove insoluble cellular debris. The supernatant was moved to a fresh 1.5mL microfuge tube and allowed to sit at room temp for at least two hours and twenty minutes before limited proteolysis (LiP), to ensure that PMSF had sufficient time to hydrolyze (approx. four half lives) and would not inhibit our protease during the LiP step. During this time, protein concentrations of the samples were determined using the bicinchoninic acid (BCA) assay, according to the manufacturer’sprotocol (Pierce Rapid Gold Protein Assay Kit, Thermo Scientific A53225). Supernatant was normalized to 0.75 mg/mL with lysis buffer. Each sample was split into si× 100μL aliquots, three for non-LiP (no PK) and three for LiP.

After adequate time for hydrolysis of PMSF has passed (so that PMSF will not inhibit PK, at least 2 h 20 min (approx. 4 half lives at pH 8.0)), LiP can be performed. Proteinase K (PK, Thermo Scientific 17916, previously prepared as 1 mg/mL stock in 1:1 v/v of lysis buffer to 20% glycerol, aliquoted, flash frozen and stored at -20°C) was diluted to 0.25 mg/mL such that 3uL of the dilution would contain 1:100 w/w of PK to the amount of sample to be digested (in this case, either 0.75μg). For each of the three LiP reactions per sample, 3μL of 0.25 mg/mL PK was placed at the bottom of a fresh 1.5mL microfuge tube. 100μL of normalized sample was added to the tube containing PK and pipetted rapidly up and down seven times to mix thoroughly. This mixture was incubated for exactly one minute before the tube was added to a 105°C mineral oil bath for five minutes to quench the PK reaction. For each sample, three corresponding no-PK, non-LiP controls were performed. In this case, 100μL of normalized lysate was added to a fresh 1.5mL tube without PK and then added to the oil bath for 5 minutes. After 5 minutes, all samples were removed from the oil bath and quickly centrifuged to collect condensation and transferred to a new 2mL tube containing 76mg urea (8M urea f.c., 314μL f.v.). To each sample in 8M urea, in order to reduce disulfides, dithiothreitol (DTT) was added to 10mM f.c. (2.25μL added from a freshly made 700mM DTT stock) and incubated in a thermomixer for 30 min at 37°C and 700 rpm. To cap cysteines, iodoacetamide (IAA) was added to a f.c. of 40mM (9μL added from a freshly made 700mM IAA stock) and incubated at room temperature (rt) in the dark for 45 min. Samples were diluted to 2M urea f.c. by adding 505μL of a freshly-made 100mM ammonium bicarbonate (ambic) stock. To each sample, 1.5μg trypsin (New England Biolabs) was added (1:50 w/w of trypsin to protein). Samples were incubated overnight (approximately 16 hours) in a thermomixer at 25°C and 700 rpm. Digested samples were acidified with trifluoroacetic acid (TFA, Acros) to a 1% f.c. by volume by adding 6.6μL TFA. Peptides were desalted using Sep-Pak Vac 1cc (50mg) C18 cartridges (Waters) and dried down and stored at -80°C, as described in section 3b.

Samples were resuspended in 75μL 0.1% formic acid and data was acquired as described in section 3c, except that samples were shot in technical duplicate for one of the three biological replicates.

Eleven label-free quantification (LFQ) analyses were performed, one with all the no-PK (trypsin only) controls for the biological replicate analysis, and one with the LiP samples (which contain PK) for the biological replicate analysis, six for each of the technical replicates (each compares the same sample that was shot twice on the instrument), and three for LiP reproducibility (one for each animal that compares the LiP performed on samples from the same animal). MSFragger and the Minora Feature Detector nodes in PD were used for the LFQs. Default settings were used except where specified otherwise. Settings were the same as in section 3e.

For the instrument variation (technical) analyses, the same sample that was injected twice was compared. For the LiP variation analyses, LiP replicates from the same sample were compared. Abundance ratios generated directly from PD were used.

For the biological replicate analysis, one of the LFQs, the control LFQ, compared the three non-PK, trypsin-only, samples for all the biological replicates. The LiP LFQ compared the three LiP samples, where PK was added. For the control LFQ, raw normalized extracted ion intensity data for the identified peptides were exported from the .pdResult file to a three-tier excel file (protein with peptide groups under each protein, with consensus features underneath each peptide group). For the LiP LFQ, ion intensity data was exported in a two-tier excel file, with peptide groups exported with consensus features underneath each peptide group. Also for the LiP LFQ, another two-tier file with peptide groups and the protein(s) they match to underneath each peptide group was exported. This two two-tier export approach was used because with three-tier files, peptides that matched to multiple proteins would be repeated underneath the other proteins it matched to.

These data were further processed using an in-house python script. Briefly, normalized ion counts were collected across the biological replicates for each peptide group. Effect sizes are the ratio of averages (reported in log2), and P-values (reported as –log10) were assessed using t tests with Welch’s correction for unequal population variances. Missing data are treated as described in section 3e, as is effect size calculation. For each peptide, the protein ratio from the control LFQ is referenced – if the protein abundance ratio is greater than two-fold in either direction, the P-value for the protein ratio is less than 0.01, and the peptide matches to a single protein, the peptide ratio is normalized by the protein abundance ratio. The script returns a file listing all the peptides that can be confidently quantified, and provides the effect sizes for all three ratios (animals B/C, A/C, and B/C).

### 9. Enzyme Activity Assays

Activity assays were performed for three AU and three AI whole hippocampi on two CASCs, phosphoglycerate kinase (PGK) and transketolase (Tkt). Tissue was Dounce homogenized on ice in 750μL buffer (100mM Tris pH 8.0, 2mM MgCl_2_) in the presence of protease inhibitors (0.5mM PMSF, 0.015mM E64, 0.05mM bestatin f.c) and let stand for 10 min. Lysate was clarified at 10000xg for 10 min at 4°C, and the supernatant was retained. Protein concentration was determined using BCA assay (Pierce Rapid Gold Protein Assay Kit, Thermo Scientific A53225), using BSA as a standard. Clarified lysate was normalized to 1.5 mg/mL in buffer (100mM Tris pH 8.0 and 2mM MgCl_2_) and kept on ice. For each activity assay, an NADH standard curve was created. Each well of the standard curve contained 0, 10, 20, 30, 40, and 50 nmol/well NADH diluted in 100mM Tris pH 8.0 (200μL f.v. each) in a transparent 96-well plate.

#### 9a. PGK Activity Assay

The PGK activity assay measures the activity of PGK by providing its substrate 3-phosphoglycerate (3-PG). PGK converts 3-PG (in the presence of excess ATP) to 1,3-bisphosphoglycerate (1,3-BPG) (this is the reverse reaction relative to the direction of glycolysis). 1,3-BPG is converted by glyceraldehyde-3-phosphate dehydrogenase (GAPDH) to glyceraldehyde-3-phosphate (GAP). This reaction consumes NADH and produces NAD^+^. NADH absorbs at 340nm, whereas NAD^+^ does not, so the disappearance of NADH can be monitored and used to determine the activity of PGK in the lysate by varying the amounts of 3-PG available for the reaction.

Each well of the reaction contains 0.5mM f.c. EDTA, 0.2mM f.c. NADH, 5mM f.c. ATP, 0.4μM f.c. GAPDH (Millipore Sigma), buffer (100mM Tris pH 8.0, 2mM MgCl_2_) and varying amounts of 3-PG. The final concentrations of 3-PG in each well is 0mM, 0.1mM, 0.25mM, 0.5mM, 1mM, 1.5mM, 2.5mM and 5mM. The final volume of all components (not including the lysate) is 100μL. The normalized lysate is diluted to 0.0375 mg/mL and 100μL of lysate is quickly added to each well (not including the NADH standard curve wells). Absorbance at 340nm was immediately measured in a microplate reader (Molecular Devices SpectraMax iD3) at 1 minute intervals for 30 minutes total.

#### 9b. Tkt Activity Assay

The tkt assay measures the activity of tkt by providing its substrate ribose-5-phosphate (R5P). Tkt converts R5P (along with xylulose-5-phosphate (X5P), which can be converted from R5P *in vivo* and is not rate-limiting) to sedoheptulose-7-phosphate and D-glyceraldehyde-3-phosphate (G3P). Triose phosphate isomerase converts G3P to dihydroxyacetone-phosphate (DHAP). DHAP is converted to glycerol-3-phosphate by α-glycerophosphate dehydrogenase (GDH), and this reaction consumes NADH and produces NAD^+^. Since NADH absorbs at 340nm, and NAD^+^ does not, disappearance of NADH can be monitored and used to determine the activity of tkt in the lysate by varying the amounts of R5P available for the reaction.

Each well of the reaction contains 2.5mM f.c. CaCl_2_, 0.4mM f.c. NADH, 80μM f.c. thiamine diphosphate (ThDP, a cofactor for tkt, Sigma-Aldrich), 1 unit per well GDH-TPI (Sigma-Aldrich), buffer (100mM Tris pH 8) and varying amounts of R5P. The final concentrations of R5P in each well are 0mM, 0.5mM, 1mM, 1.5mM, 2mM, 3mM, 5mM, and 8mM. The final volume of all components (not including the lysate) is 100μL. 100μL of 1.5 mg/mL lysate is quickly added to all sample wells (not including NADH standard curve wells) and absorbance is immediately measured at 340nm for 30 min at 1 min intervals on a microplate reader (Molecular Devices SpectraMax iD3).

#### 9c. Activity Assay Data Analysis

Reaction rate was calculated as the slope of the linear region of the absorbance curve. Corrected reaction rates were obtained by subtracting the rate of the no-substrate reaction from all other rates. Since a disappearance of substrate was measured, the absolute value of the rates was used. Michaelis-Menten best-fit values for K_m_ and V_max_ were obtained by performing a Michaelis-Menten nonlinear fit of substrate concentration versus rate, using Graphpad Prism version 10.

### 10. Western Blotting

Hippocampal lysate prepared for activity assays was diluted with water and 5X Tris-glycine-SDS loading buffer such that a 15μL aliquot contains 5μg of lysate and 1X loading buffer. Samples were boiled for 10 min at 90°C, and quickly spun down to collect condensation. 15μL of each sample and 2μL PageRuler prestained plus ladder (Thermo Fisher) was loaded into a precast Novex WedgeWell 4-20% Tris-Glycine protein gel (Invitrogen). The protein gel was run at 200V for approx. 40 min. After electrophoresis, the gel was transferred to a PVDF membrane (iBlot 2 PVDF mini stack, Invitrogen) using an iBlot 2 gel transfer device (Invitrogen) at 20V for 7 min. Membrane was cut as appropriate and blocked for 2 hours in 10mL 5% (w/v) nonfat milk in TBST (20mM Tris pH 7.5, 150mM NaCl, 0.1% Tween-20, with tween added the same day as used). The blocked membrane was incubated approx. 16 hours with rocking with the appropriate primary antibody at the appropriate dilution: rabbit anti-PGK (1:5000 dilution, Invitrogen MA5-32174), rabbit anti-transketolase (1:10000 dilution, Proteintech 11039-1-AP), and rabbit anti-alpha/beta tubulin (1:5000 dilution, Cell Signaling Technology 2148S). Membranes were rinsed with rocking in 10mL TBST for 10 min, three times. A goat anti-rabbit horseradish peroxidase (HRP) secondary antibody (1:10000 dilution, Invitrogen 65-6120) was added to 5% (w/v) nonfat milk in 10mL TBST and incubated with rocking for 2 hours. The membranes were rinsed with rocking in 10mL TBST for 10 min, three times. Membranes were incubated for 5 min in 8mL f.v. of reagents from the SuperSignal West Pico Plus Chemiluminescent Substrate kit (Thermo Fisher). Membranes were imaged on a ChemiDoc Touch Imaging System (Bio-Rad). Images were analyzed by calculating integrated density using ImageJ.

## Acknowledgments

H.E.T. acknowledges support from the Chemistry-Biology Interface NIH training grant program (T32-GM080189 and T32-GM149382). H.E.T. acknowledges support from the age-related neuroscience NIH training grant (T32-AG027668). S.D.F. acknowledges support from the NIH Director’s New Innovator Award (DP2-GM140926), from the National Science Foundation (MCB-2045844), from a Camille Dreyfus Teacher-Scholar Award, and from a Sloan Fellowship. S.D.F. acknowledges a Longevity Impetus Grant from Norn Group, Hevolution Foundation and Rosenkranz Foundation. The authors thank Robert McMahan, Jala Atufa, and Ashley Becker for assistance with MWM behavioral characterization and hippocampal dissections. The authors thank Michela Gallagher for directing the animal resource, insightful conversations, and critical review of the manuscript.

## Funding

National Science Foundation MCB-2045844 (SDF)

National Institutes of Health DP2-GM140926 (SDF)

National Institutes of Health T32-GM149382 (HET)

National Institutes of Health T32-AG027668 (HET)

Camille Dreyfus Teacher-Scholar Award (SDF)

Sloan Fellowship (SDF)

Longevity Impetus Grant (SDF)

National Institutes of Aging P01AG009973

## Author contributions

Conceptualization: HET, SDF

Methodology: HET, AB, SDF

Investigation: HET, AB

Visualization: HET, SDF

Supervision: SDF

Writing—original draft: HET, SDF

Writing—review & editing: HET, AB, SDF

## Competing interests

The authors declare that they have no competing interests.

## Data and materials availability

The mass spectrometry proteomics data have been deposited to the ProteomeXchange Consortium via the PRIDE partner repository with the dataset identifier PXD053728 and PXD052770. Peptide quantifications and code used to analyze the LiP-MS data are available on Zenodo at <u>10.5281/zenodo.13795773</u>. Protein summary files are available as excel supplemental datasets.

## Supplemental Figures

**Figure S1.**
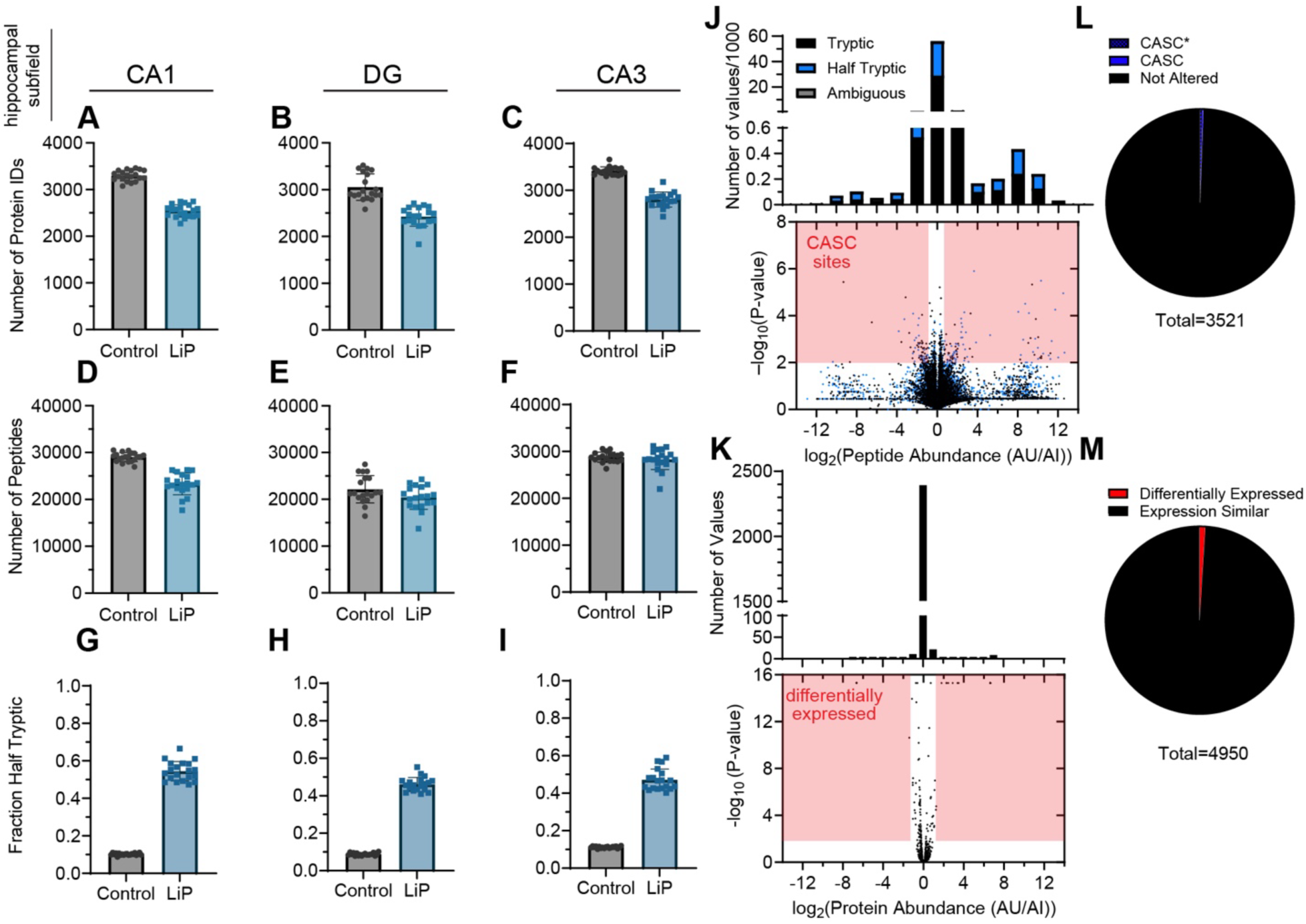
Quality Controls for Limited Proteolysis Mass Spectrometric (LiP-MS) Study of the Aging Rat’s Hippocampal Proteome. (A-C) Number of proteins identified in individual mass spec runs on three hippocampal subfields ((A) CA1; (B) DG; (C) CA3) filtered to an FDR of 5%. Each point represents an individual subject (biological replicate). Bars represent average, errors bars represent standard deviations. (D-F) Number of peptides identified in individual mass spec runs on three hippocampal subfields ((D) CA1; (E) DG; (F) CA3) filtered to an FDR of 5%. Each point represents an individual subject (biological replicate). Bars represent average, errors bars represent standard deviations. (G-I) Of the peptides identified in an individual mass spec run on three hippocampal subfields ((G) CA1; (H) DG; (I) CA3), the fraction that are half-tryptic (that is, one cut-site is non-tryptic and presumed to arise from Proteinase K). Bars represent average, errors bars represent standard deviations. In panels (A)-(I), “Control” denotes samples processed without limited proteolysis (only digested with trypsin) and “LiP” denotes samples subjected to limited proteolysis with Proteinase K, then trypsin digest. (J) Volcano plot showing changes in peptide abundance of tryptic (black) and half-tryptic (blue) peptides in the CA3 subfield between AU and AI rats. Dots that fall in the regions in red are deemed significant based on effect size (>2-fold) and p < 0.01 by t-test with Welch’s correction for unequal population variance. (K) Volcano plot showing changes in protein abundance in the CA3 subfield between AU and AI rats. Dots that fall in the regions in red are deemed significant based on effect size (>2-fold) and p < 0.01 by t-test with Welch’s correction for unequal population variance. (L) Number of proteins with cognition-associated structural changes (CASC) in the CA3 subfield. CASC proteins have two or more peptides with significant changes. CASC* denotes proteins in which one (or both) of the significant peptides could be assigned to multiple isoforms. (M) Number of proteins with different measured abundance in AU vs AI subjects in the CA3 subfield.

**Figure S2.**
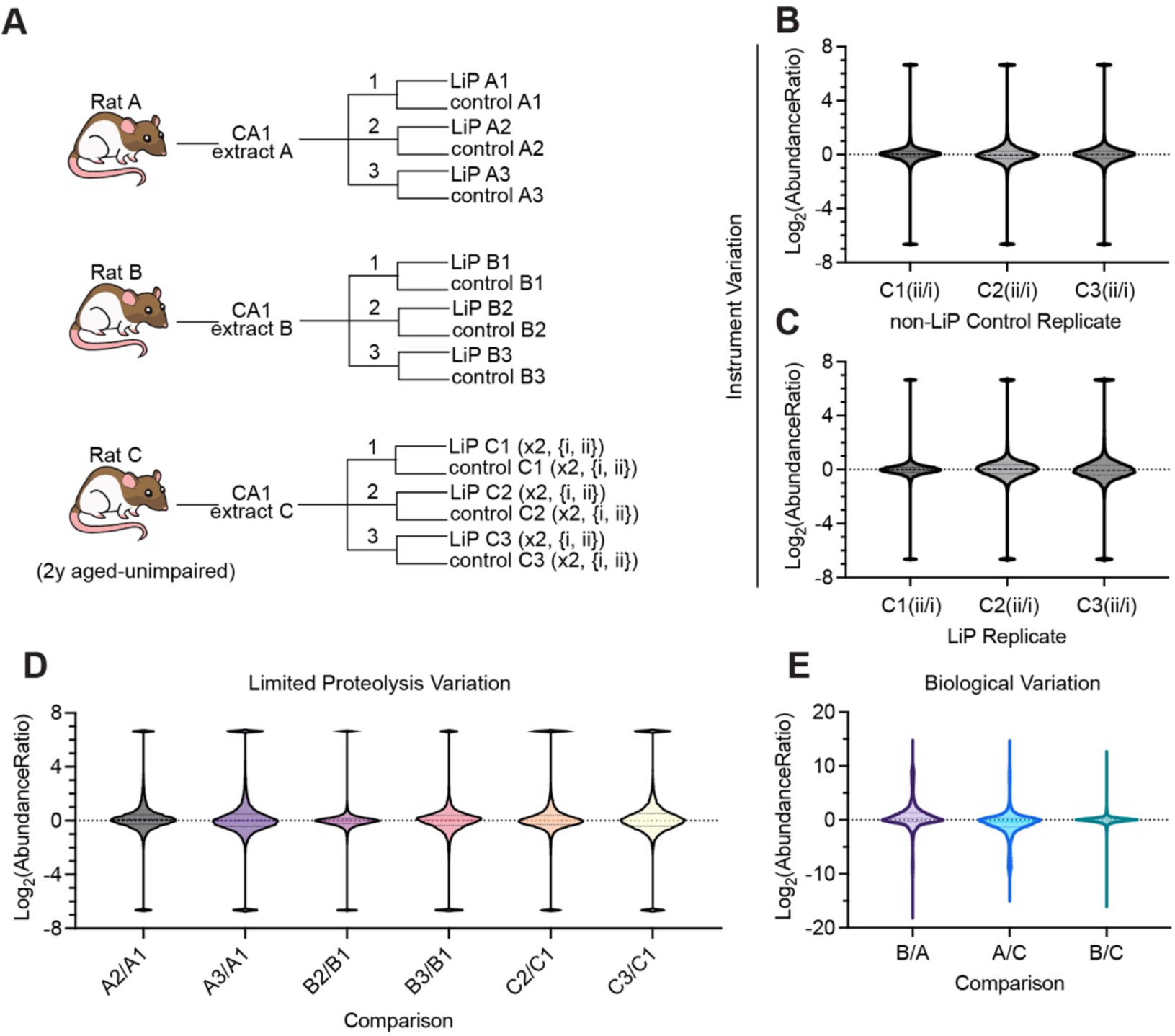
Reproducibility of Limited Proteolysis Methodology. (A) Scheme illustrating experimental design of the reproducibility study. Three biological replicates (all cognitively unimpaired rats, aged 2 y) were sacrificed, and the CA1 subfield of their hippocampi extracted. These biological replicates were subjected to 3 technical replicates of preparing LiP samples and “control” (non-LiP trypsin-only) samples. For rat C, these 3 technical replicates were also subjected to analytical duplicates (injected on the LC-MS/MS twice). (B, C) Analytical variability in peptide abundance between two injections of non-LiP control samples, (B); and between two injections of LiP samples, (C). (D) Technical variability in peptide abundance following replicate LiP reactions derived from the same biological replicate. (E) Biological variability in peptide abundance across different subjects’ CA1 hippocampal region. For each subject, peptide abundance signals were averaged across the three technical replicates.

**Figure S3.**
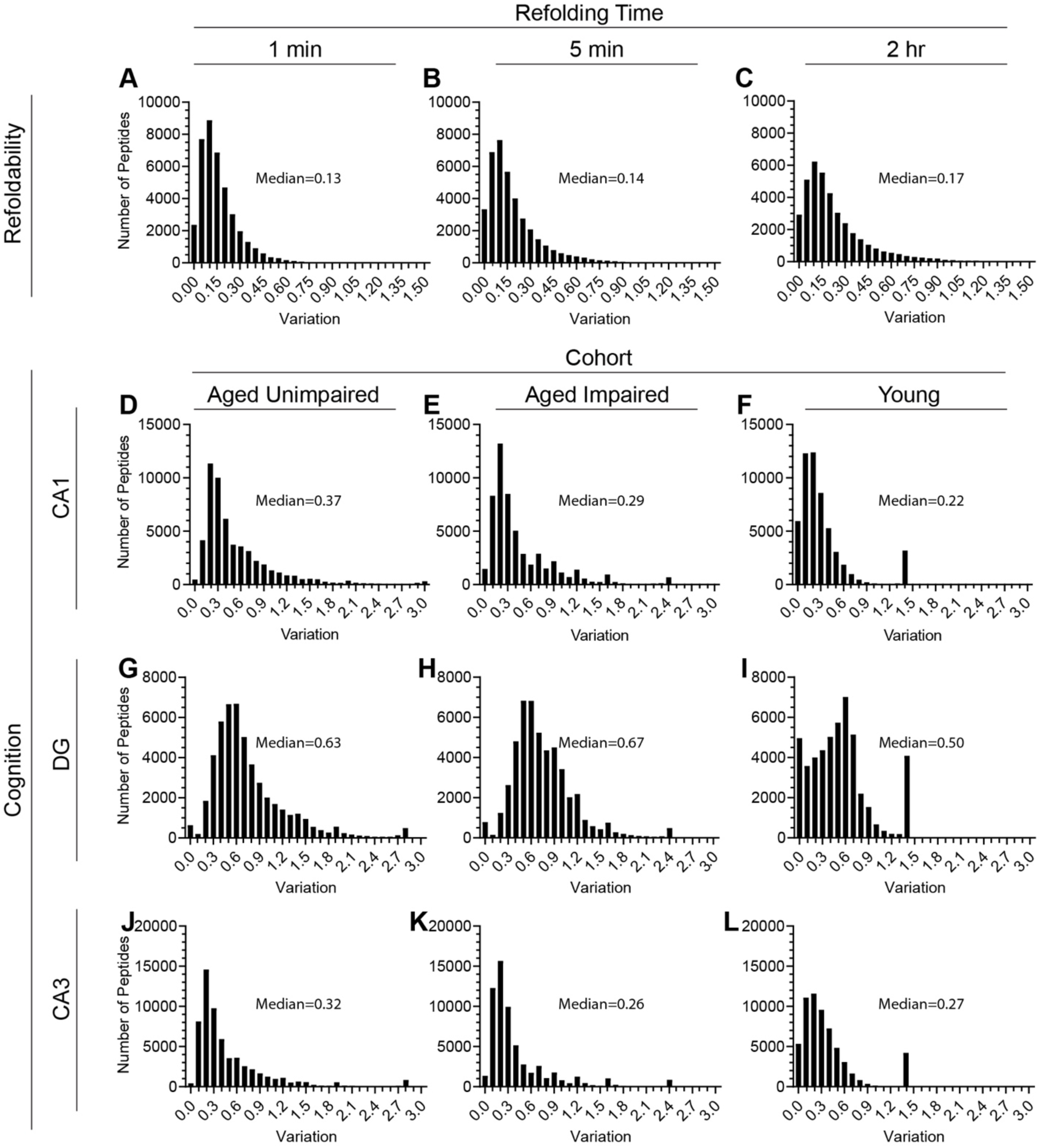
Analysis of Variation. Histograms show the coefficients of variation (standard deviation divided by mean) for peptide abundance in limited proteolysis samples across the replicates for all conditions surveyed in this study. Median coefficient of variation reported. (A-C) Coefficients of variation across 3 replicates of global refolding reactions, diluted from denaturant and allowed to refold for (A) 1 min; (B) 5 min; (C) 2 h. (D-L) Coefficients of variation across biological replicates from 3 hippocampal subfields (CA1, panels (D-F); DG, panels (G-I); CA3, panels (J-L)) collected from rats from 3 cohorts (aged unimpaired, panel (D), 10 replicates, panels (G),(J), 9 replicates; aged impaired, panels (E),(H),(K), 7 replicates; young, panels (F),(I),(L), 3 replicates).

**Figure S4.**
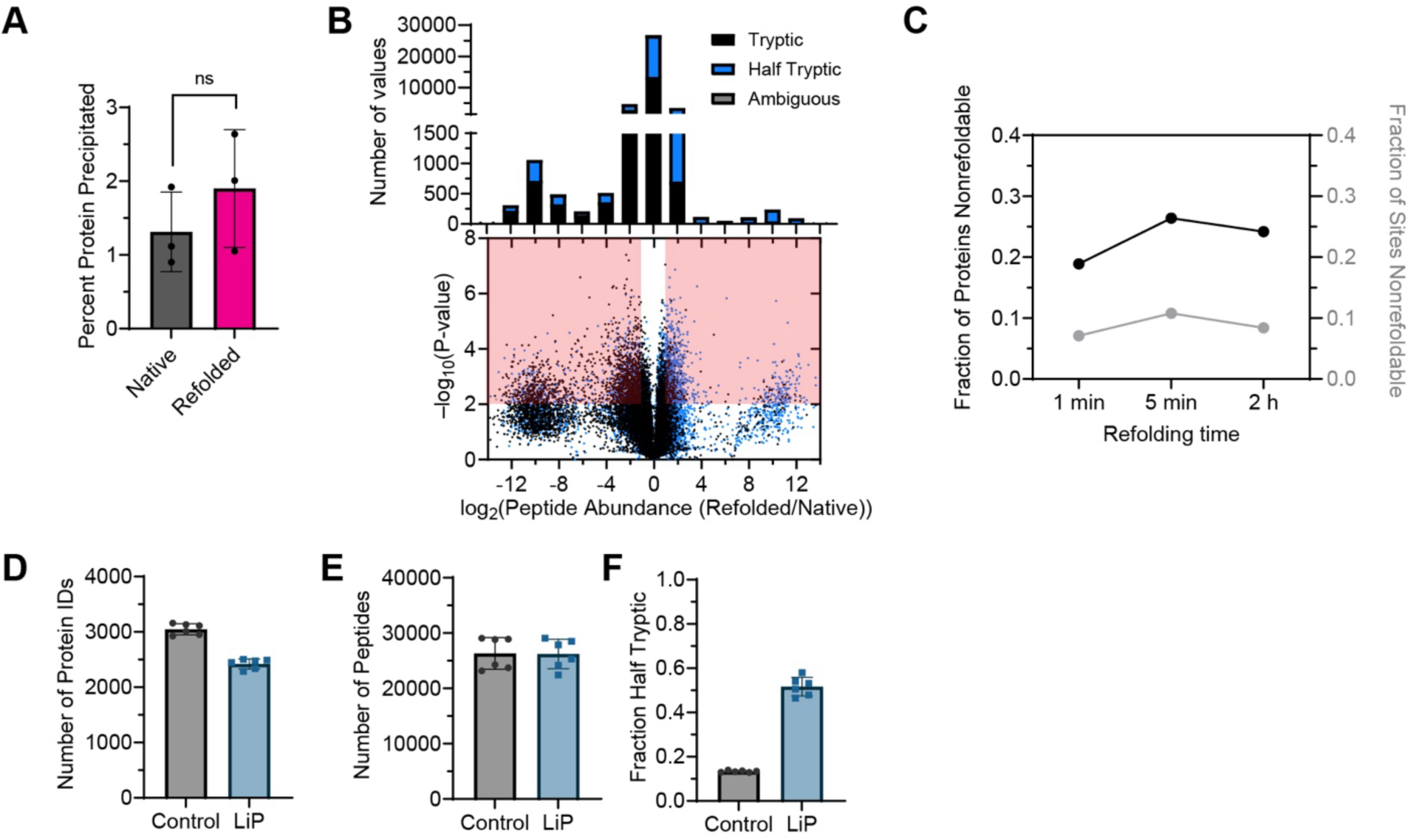
Quality Controls for Limited Proteolysis Mass Spectrometric (LiP-MS) Refolding Experiment on Rat Hippocampal Proteome. (A) Percent of protein content that precipitates from clarified rat hippocampal extract either before (gray) or after (pink) a global refolding cycle conducted by incubating in 6 M guanidinium chloride followed by 100-fold dilution (see *Methods* Section 4 for details). Aggregated protein was collected by centrifugation, resuspended in 8 M urea, and quantified by bicinchoninic acid (BCA) assay. Dots represent biological replicates, bars represent averages, and error bars represent standard deviations. (B) Volcano plot showing changes in peptide abundance of tryptic (black) and half-tryptic (blue) peptides in the rat hippocampus following a global unfolding-refolding cycle relative to a native reference using a 5 min refolding time. Dots that fall in the regions in red are deemed significant based on effect size (>2-fold) and p < 0.01 by t-test with Welch’s correction for unequal population variance. Histograms show number of peptides possessing abundance ratios in the various ranges. (C) Fraction of proteins nonrefoldable (black, using the criterion that the protein has two or more peptides with significant changes in proteolytic susceptibility in the refolding reactions) and fraction of peptides/sites nonrefoldable (gray, using the criterion the peptide has >2-fold change in abundance (p < 0.01 by t-test with Welch’s correction) in the refolding reactions) as a function of refolding time. Each time-point used three separate refolding reactions performed on biological triplicate (i.e., separate hippocampi from separate rats). (D) Number of proteins identified in individual mass spec runs used for *in vitro* refolding experiments filtered to an FDR of 5%. Bars represent average, errors bars represent standard deviations. (E) Number of peptides identified in individual mass spec used for *in vitro* refolding experiments filtered to an FDR of 5%. Bars represent average, errors bars represent standard deviations. (F) Of the peptides identified in an individual mass spec run used for *in vitro* refolding experiments, the fraction that are half-tryptic (that is, one cut-site is non-tryptic and presumed to arise from Proteinase K). Bars represent average, errors bars represent standard deviations. In panels (D)-(F), “Control” denotes samples processed without limited proteolysis (only digested with trypsin) and “LiP” denotes samples subjected to limited proteolysis with Proteinase K, then trypsin digest.

**Figure S5.**
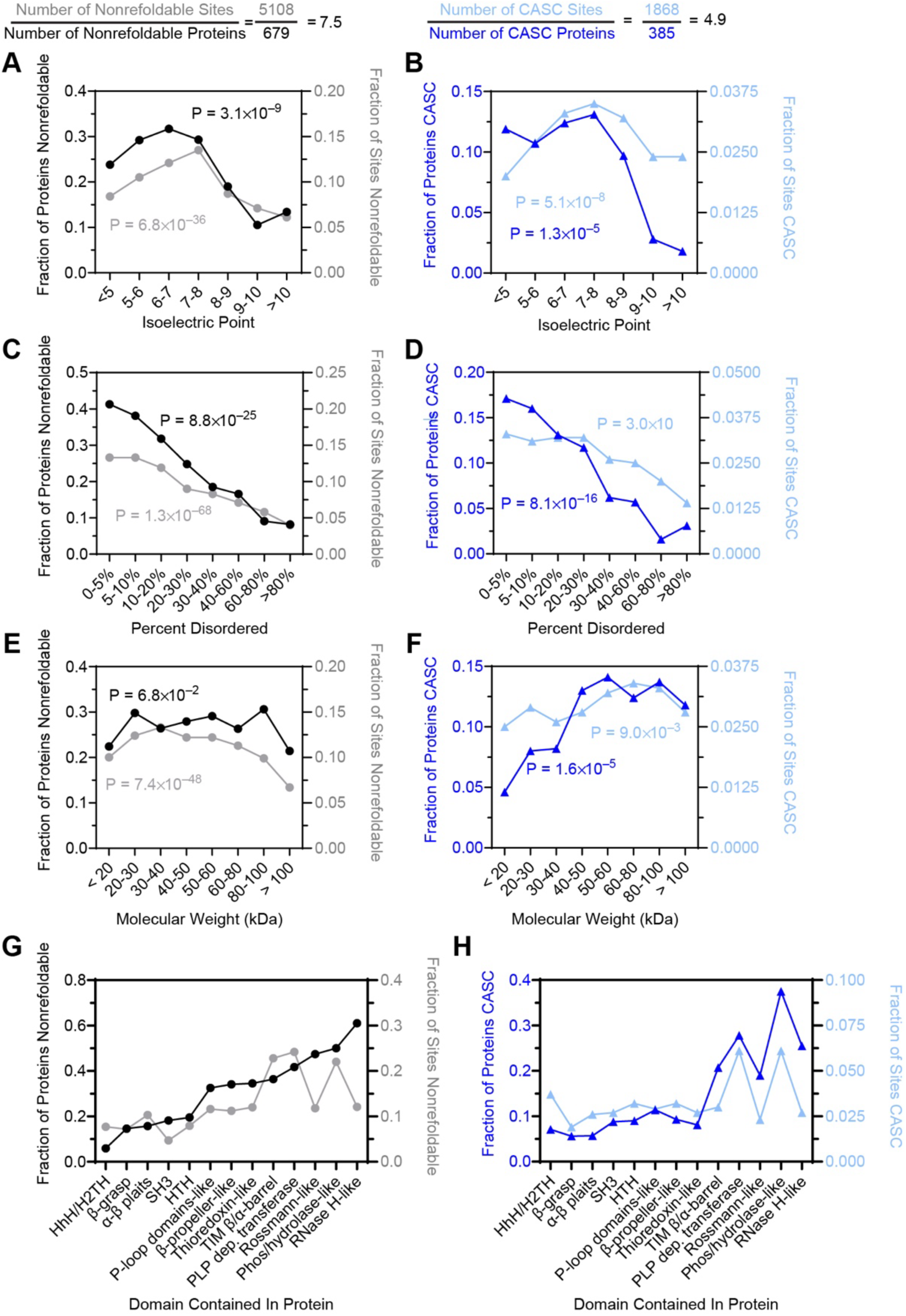
Robustness Analysis of Trends Found on Nonrefoldable Proteins and Cognition-Associated Structurally Changed (CASC) Proteins. (A,C,E,G) Plots provide the fraction of proteins within a given category that are nonrefoldable after allowed 5 min to refold from denaturant (black, left y-axis) and the proportion of sites (peptides) within a given category that had significantly altered proteolytic susceptibility after 5 min of refolding (gray, right y-axis). Proteins and peptides are divided as a function of: (A) protein isoelectric point (pI), (C) percent disorder according to Metapredict, and (E) molecular weight. P-values (according to chi-square test) against the null hypothesis that nonrefoldability is independent of the categorical variable in question are provided. (G) Proteins (or peptides) assessed based on whether they contain a domain of a given topology (or come from a protein that contains a domain of a given topology). These categories are not mutually exclusive since some proteins contain multiple domains. (B,D,F,H) Plots provide the fraction of proteins within a given category that are CASC in the CA1 region (blue, left y-axis) and the proportion of sites (peptides) within a given category that had significantly altered proteolytic susceptibility in aged impaired rats compared to aged unimpaired (light blue, right y-axis). Proteins and peptides are divided as a function of: (B) protein isoelectric point (pI), (D) percent disorder according to Metapredict, and (F) molecular weight. P-values (according to chi-square test) against the null hypothesis that CASC is independent of the categorical variable in question are provided. (H) Proteins (or peptides) assessed based on whether they contain a domain of a given topology (or come from a protein that contains a domain of a given topology). These categories are not mutually exclusive since some proteins contain multiple domains.

**Figure S6.**
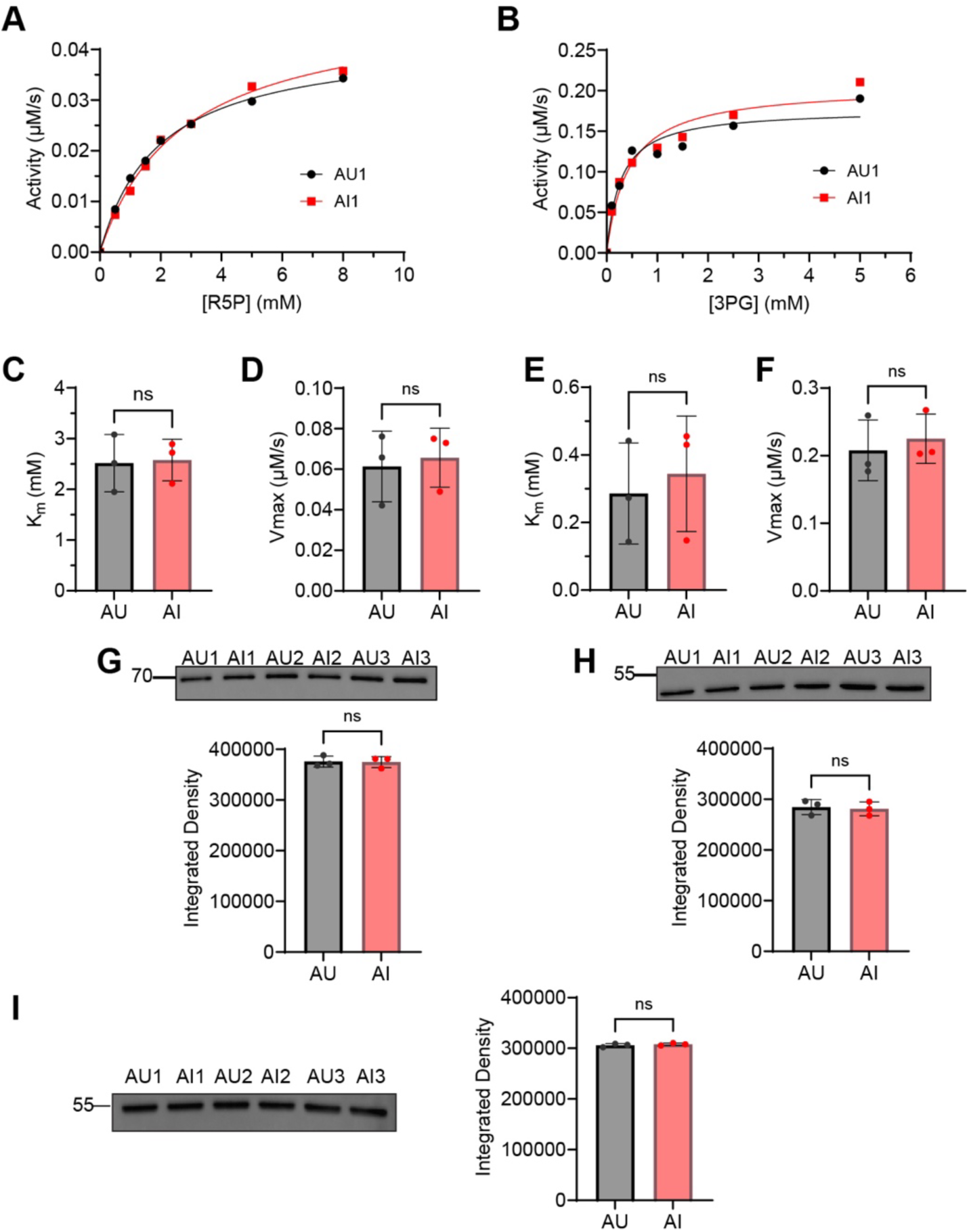
Activity Assays. (A) Representative Michaelis-Menten kinetics on rat hippocampal transketolase, measured in lysate, by titrating the substrate D-ribose-5-phosphate (R5P), see *Methods* for assay details. Shown is activity measured from one biological replicate of an aged impaired (AI) rat and one biological replicate of an aged unimpaired (AU) rat, at distinct R5P concentrations. (B) Representative Michaelis-Menten kinetics for the reverse reaction of phosphoglycerate kinase (PGK) from rat hippocampi, measured in lysate, by titrating the product 3-phosphoglycerate (3PG), see *Methods* for assay details. Shown is activity measured from one biological replicate of an aged impaired (AI) rat and one biological replicate of an aged unimpaired (AU) rat, at distinct 3PG concentrations. (C, D) Fit parameters to *K*_m_ (C) and *V*_max_ (D) from three biological replicates for transketolase. (E, F) Fit parameters to *K*_m_ (E) and *V*_max_ (F) from three biological replicates for PGK. (G) Western Blot against rat transketolase from same 6 extracts used for kinetics measurements and bar chart showing band quantifications by densitometry in ImageJ. Near equal transketolase abundances suggests *V*_max_’s are comparable without normalization. (H) Western Blot against rat PGK from same 6 extracts used for kinetics measurements and bar chart showing band quantifications by densitometry in ImageJ. Near equal transketolase abundances suggests *V*_max_’s are comparable without normalization. (I) Western Blot against rat β-tubulin from same 6 extracts used in kinetics and other Western blots, used as loading control.

**Figure S7.**
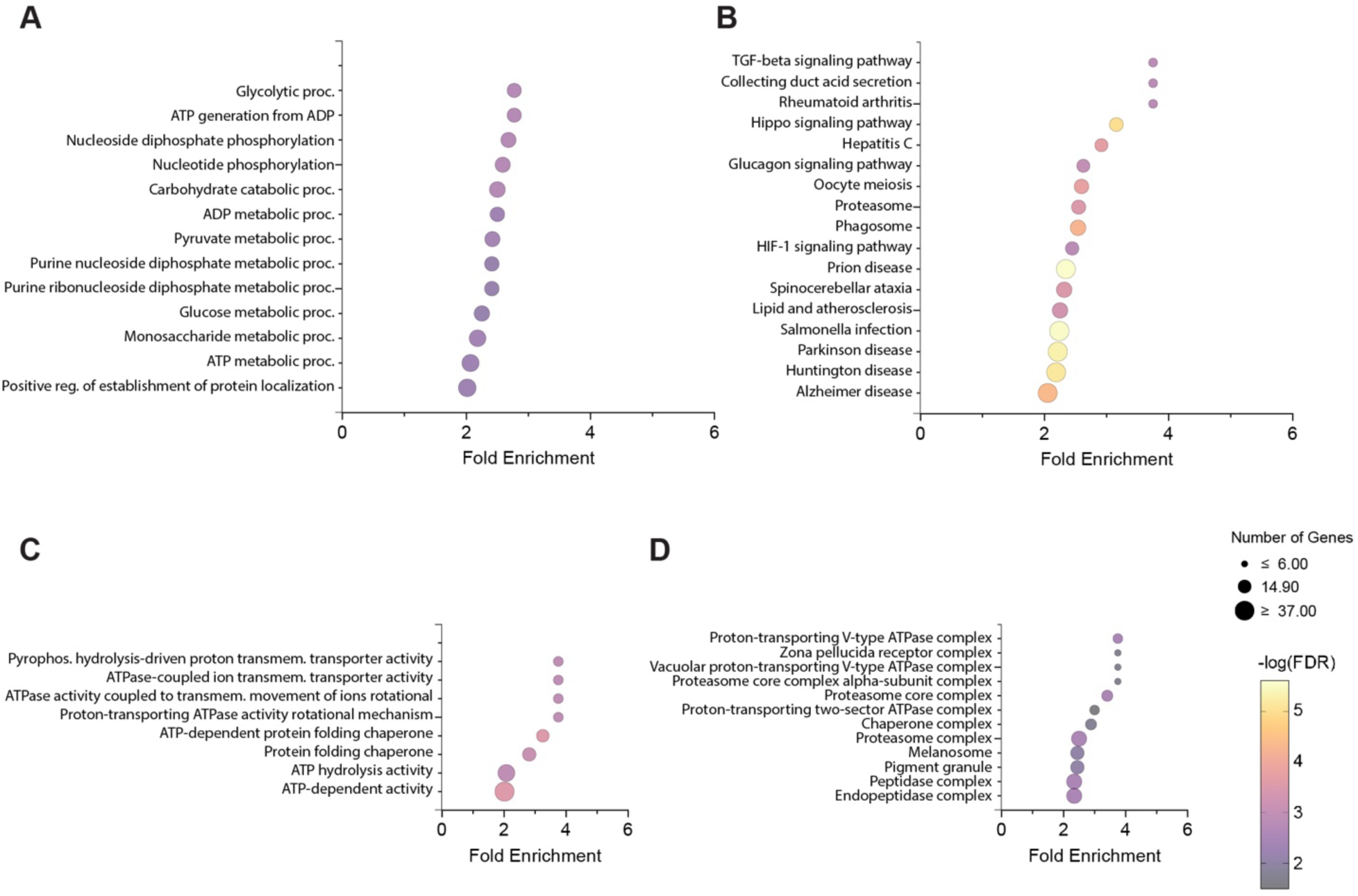
Gene Ontology Analysis of Nonrefoldable Proteins. Analysis is based on the 679 nonrefoldable proteins identified in global refolding experiments on the rat hippocampal proteome, with a reference set composed of all 2573 proteins quantified with 2 or more peptides across the 6 samples used for the experiment (biological triplicates of native and refolded (5 min) extracts) using ShinyGO (*56*). (A) Using the biological process GO terms. (B) Using the Kyoto Encyclopedia of Genes and Genomes (KEGG) database. (C) Using the molecular function GO terms. (D) Using cellular component GO terms.

**Figure S8.**
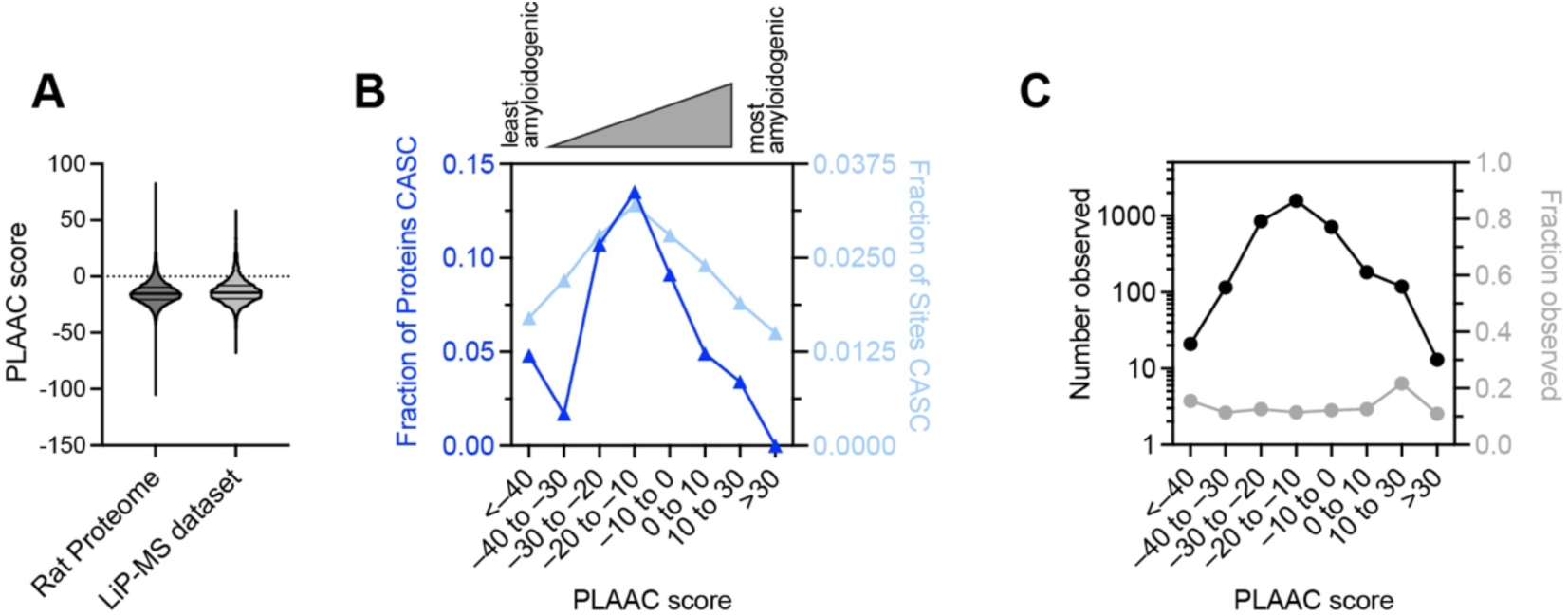
Association between CASC proteins and amyloidogenicity. Amyloid propensity is based on the log-likelihood score of the PLAAC algorithm (*72*) using default parameters. (A) Distribution of PLAAC scores for all proteins in the rat proteome versus those represented in the LiP-MS aging study presented here. (B) Plot provides the fraction of proteins that are CASC in the CA1 region (blue, left y-axis) and the proportion of sites (peptides) that had significantly altered proteolytic susceptibility in aged impaired rats compared to aged unimpaired (light blue, right y-axis), divided by each protein’s PLAAC log-likelihood score. The trend is non-monotonic, and proteins that are predicted to be most amyloidogenic (high positive PLAAC score) are less likely to be CASC. (C) Number of proteins observed in each of the PLAAC score bins (black) in the LiP-MS aging study, and the percentage of all rat proteins within a PLAAC score bin that are observed in the LiP-MS aging study (gray). This shows that although there are, for instance, many fewer proteins with PLAAC > 30, this category is nevertheless observed with the same frequency as other categories.

**Figure S9.**
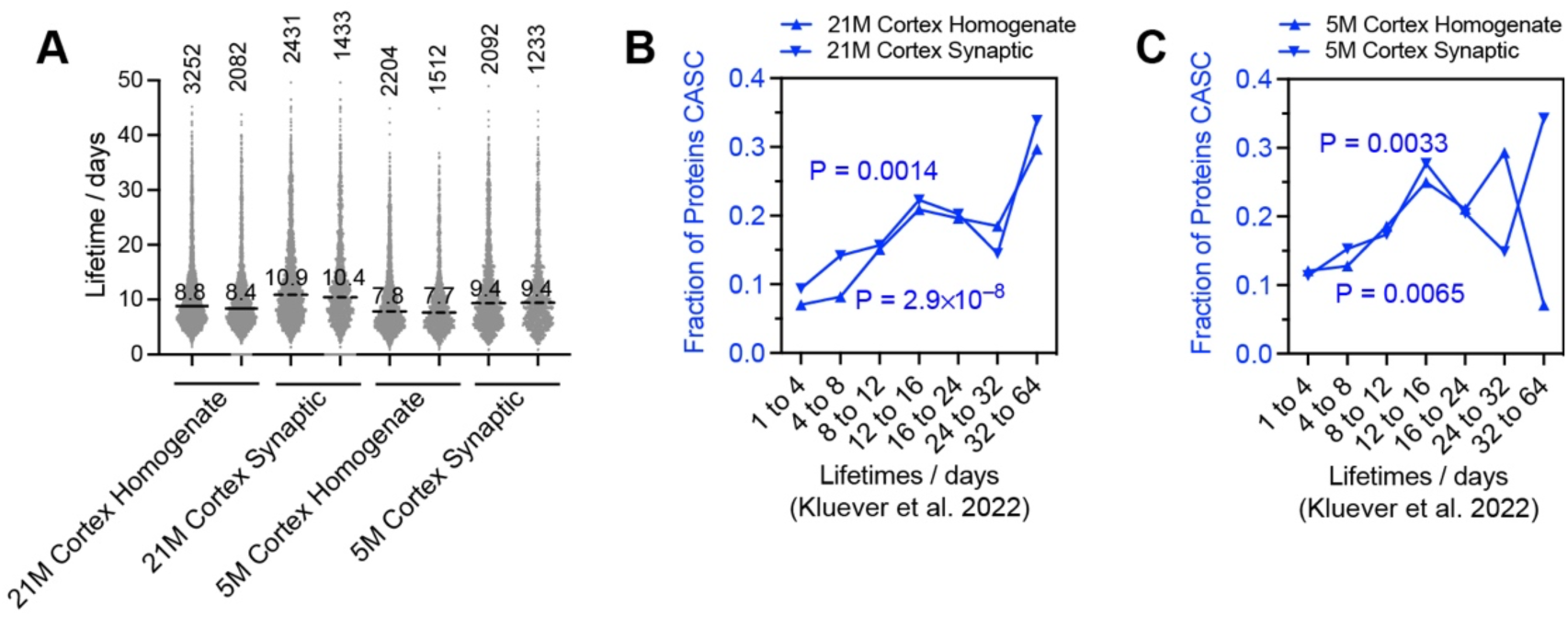
Association between CASC proteins and lifetime. Lifetimes are based on the study by Kluever et al., by referring to the values in “tHalf” columns in Table S1 and Table S3 from ref. (*79*). As the previous work used mice instead of rats, we used InParanoiDB 9 to identify the rat orthologue for each mouse protein. (A) Distribution of protein lifetimes observed by Kluever et al. for proteins isolated from mouse cortex, either from animals aged to 21 months (old) or 5 months (young), or from total cortex homogenates or synaptic fractions. To the right of each, the distribution of protein lifetimes from that experiment that were also assessed in the LiP-MS aging study. The median lifetime and the total number of proteins measured are noted for each distribution. (B) Plot provides the fraction of proteins that are CASC in the CA1 region (blue, left y-axis), divided by the orthologous protein’s lifetime in the mouse cortex. The trend is non-monotonic, and proteins that are predicted to be most amyloidogenic (high positive PLAAC score) are less likely to be CASC. P-values (according to chi-square test) against the null hypothesis that CASC is independent of the lifetime category. (C) As panel B, except using lifetimes measued for rats aged to 5 months instead of 21 months.

## Notes

https://zenodo.org/records/13795774

### Competing Interest Statement

The authors have declared no competing interest.

